# Closed-loop auditory stimulation method to modulate sleep slow waves and motor learning performance in rats

**DOI:** 10.1101/2021.03.17.435881

**Authors:** Carlos G. Moreira, Christian R. Baumann, Maurizio Scandella, Sergio I. Nemirovsky, Sven Leach, Reto Huber, Daniela Noain

**Affiliations:** Department of Neurology, University Hospital Zurich, University of Zurich, Zurich, Switzerland; University Center of Competence Sleep & Health Zurich (CRPP), University of Zurich, Zurich, Switzerland; Neuroscience Center Zurich (ZNZ), Zurich, Switzerland; Institute of Biological Chemistry, School of Exact and Natural Sciences (IQUIBICEN). CONICET – University of Buenos Aires, Buenos Aires, Argentina; Child Development Center, University Children’s Hospital Zurich, Zurich, Switzerland; Department of Child and Adolescent Psychiatry and Psychotherapy, Psychiatric Hospital, University of Zurich

**Keywords:** Closed-loop auditory stimulation, sleep slow waves, motor learning, sleep modulation

## Abstract

Slow waves and cognitive output have been modulated in humans by phase-targeted auditory stimulation. However, to advance its technical development and further our understanding, implementation of the method in animal models is indispensable. Here, we report the successful employment of slow waves’ phase-targeted closed-loop auditory stimulation (CLAS) in rats. To validate this new tool both conceptually and functionally, we tested the effects of up- and down-phase CLAS on proportions and spectral characteristics of sleep, and on learning performance in the single pellet-reaching task, respectively. Without affecting 24-h sleep-wake behavior, CLAS specifically altered delta (slow waves) and sigma (sleep spindles) power persistently over chronic periods of stimulation. Down-phase CLAS exerted a detrimental effect on overall engagement and success rate in the behavioral test, and overall CLAS-dependent spectral changes were positively correlated with learning performance. Altogether, our results provide proof-of-principle evidence that phase-targeted CLAS of slow waves in rodents is efficient, safe and stable over chronic experimental periods, enabling the use of this high-specificity tool for basic and preclinical translational sleep research.

## Introduction

Non-invasive neuromodulation strategies are en vogue for their unique potential to diagnose and treat neurological and psychiatric disorders, or to rebalance the activity in dysfunctional brain networks. In particular, modulation of slow wave sleep (SWS) has increasingly gained attention over recent years. Tailoring novel stimulation tools to effectively optimize SWS profiles appears crucial to intervene SWS-related cognition (Mander et al., 2013; Papalambros et al., 2017), as well as other important brain and body processes, in both rodents and humans. Therefore, innovative approaches to further comprehend and eventually enable therapeutic implementations of SWS modulation have been tested (Marshall, Helgadóttir, Mölle, & Born, 2006; Massimini et al., 2007; Vyazovskiy, Faraguna, Cirelli, & Tononi, 2009). More recently, auditory stimulation during SWS was successfully implemented in human subjects in laboratory-based settings. At first, the method disregarded the phase of the ongoing oscillatory activity in the brain (H. V. Ngo, Claussen, Born, & Mölle, 2013; Tononi, Riedner, Hulse, Ferrarelli, & Sarasso, 2010), but was later developed further to deliver auditory stimulation in synchrony with the brain’s own rhythm in a closed-loop manner (H.-Viet V. Ngo, Martinetz, Born, & Mölle, 2013). Targeting ongoing slow waves in their up-phase enhanced slow oscillations during SWS, while targeting the waves’ down-phase had the opposite effect. The success of the human implementation of CLAS additionally encouraged the development of portable devices enabling acoustic stimulation in a home-based environment (Ferster, Lustenberger, & Karlen, 2019).

Memory formation is one of the most intriguing sleep-dependent brain processes for which the links between SWS — characterized by brain slow oscillatory activity in the delta frequency-range (0.5 – 4 Hz) and oscillations <1 Hz among its most distinctive features in the human electroencephalogram (EEG) (Timofeev, 2011) — and motor learning (Fischer, Hallschmid, Elsner, & Born, 2002; Stickgold, 2005; Walker, Brakefield, Morgan, Hobson, & Stickgold, 2002), synaptic downscaling (Tononi & Cirelli, 2014), and consolidation mechanisms (Diekelmann & Born, 2010; Rasch & Born, 2013) have been exhaustively studied. In this context, it was not surprising that the novel specific auditory modulation of SWS methods implemented in humans were first assessed in their capacity to interact with memory performance. Initially, Ngo and colleagues found enhanced performance during a paired-associates learning task in subjects undergoing up-phase auditory stimulation (H.-Viet V. Ngo et al., 2013), and soon several other studies employing similar stimulation paradigms replicated the enhanced memory effect, although with more modest effect sizes (Leminen et al., 2017; Ong et al., 2016; Papalambros et al., 2017). In a novel implementation of this concept, some of us found that auditory disruption of slow waves perturbing local SWS in the motor cortex attenuated the brain’s SWS-dependent capacity to undergo neuroplastic changes (Fattinger et al., 2017), corroborating that SWS integrity is critical to maintain cognitive efficiency. However, some authors proposed that enhancing slow waves and spindles by auditory stimulation was insufficient to improve memory above sham (Cox, Hofman, de Boer, & Talamini, 2014; H. V. Ngo, Seibold, Boche, Mölle, & Born, 2019; Weigenand, Mölle, Werner, Martinetz, & Marshall, 2016), suggesting that the effects of auditory stimulation on memory are moderate and highly variable inter and intra-studies, and evidencing that a better understanding of the method is needed.

Therefore, although a highly promising tool, the precise effects exerted by phase-targeted auditory stimulation at both electrophysiological and behavioral levels are not understood well enough yet. Moreover, effects of long-term interventions, as well as parameters optimization remain to be addressed. These limitations can partly be attributed to the lack of a closed-loop auditory stimulation (CLAS) paradigm in animals to facilitate both basic and preclinical research. For the first time, we explored here the effects exerted by phase-targeted CLAS of slow waves on slow wave activity (SWA) in healthy rats. We conceptualized that up-phase CLAS will boost delta power during non-rapid eye movement (NREM) sleep, the sleep behavioral state rich in slow waves in animals, while down-phase CLAS will have the opposite effect. As a functional readout of CLAS of slow waves, we evaluated learning performance in the single-pellet reaching task.

## Results

### CLAS’ precision and phase detection

To study the effect of CLAS of slow waves in healthy rats (**Fig. 1 and Fig. 1-figure supplement 1**), we recorded animals during an undisturbed 24-h baseline (BL) period and, thereafter, delivering up-phase, mock or down-phase auditory stimulation continuously for 16 days, while undergoing behavioural training in the single pellet reaching task (SPRT) (**Fig. 2**). Online NREM sleep staging is a crucial step during CLAS of slow waves. Briefly, our online NREM sleep staging tool consists of three real-time decision features: root mean square (rms) of delta and beta frequency bands, and EMG. In a separate cohort (14 animals: up-phase (n = 5), mock (n = 4) or down-phase (n = 5)), we assessed the performance of our online NREM sleep staging process, as well as the accuracy of our phase-detector. Overall, online NREM sleep staging had a mean sensitivity of 12% and specificity of 98% (**Fig. 3a**). Regarding online precision of stimulation in NREM sleep, 70% of epochs were confirmed offline as NREM sleep, whereas 18% were later identified as wakefulness and 2% as REM sleep (**Fig. 3b**). Phase histograms showed high accuracy in trigger distribution relative to the intended targets of 65° and 270° for up- and down-phase stimulation, respectively. Although different phases were targeted in this cohort, the phase-detector is, in its core and performance, the same. Up-phase paradigm revealed *M* = 74.97° (*SD* = 38.10°; *Var* = 12.67°) while down-phase stimulation indicated a mean target *M* = 280.55° (*SD* = 49.67°; *Var* = 19.75°). In mock animals, we confirmed onset of muted-triggers around the positive peak of the slow wave (*M* = 101.20°; *SD* = 46.89°; *Var* = 19.19°) (**Fig. 3c**). The difference in variances reflect the gradual loss of accuracy when higher phases are being targeted, an intrinsic characteristic of our phase-detection feature.

**Figure 1.**
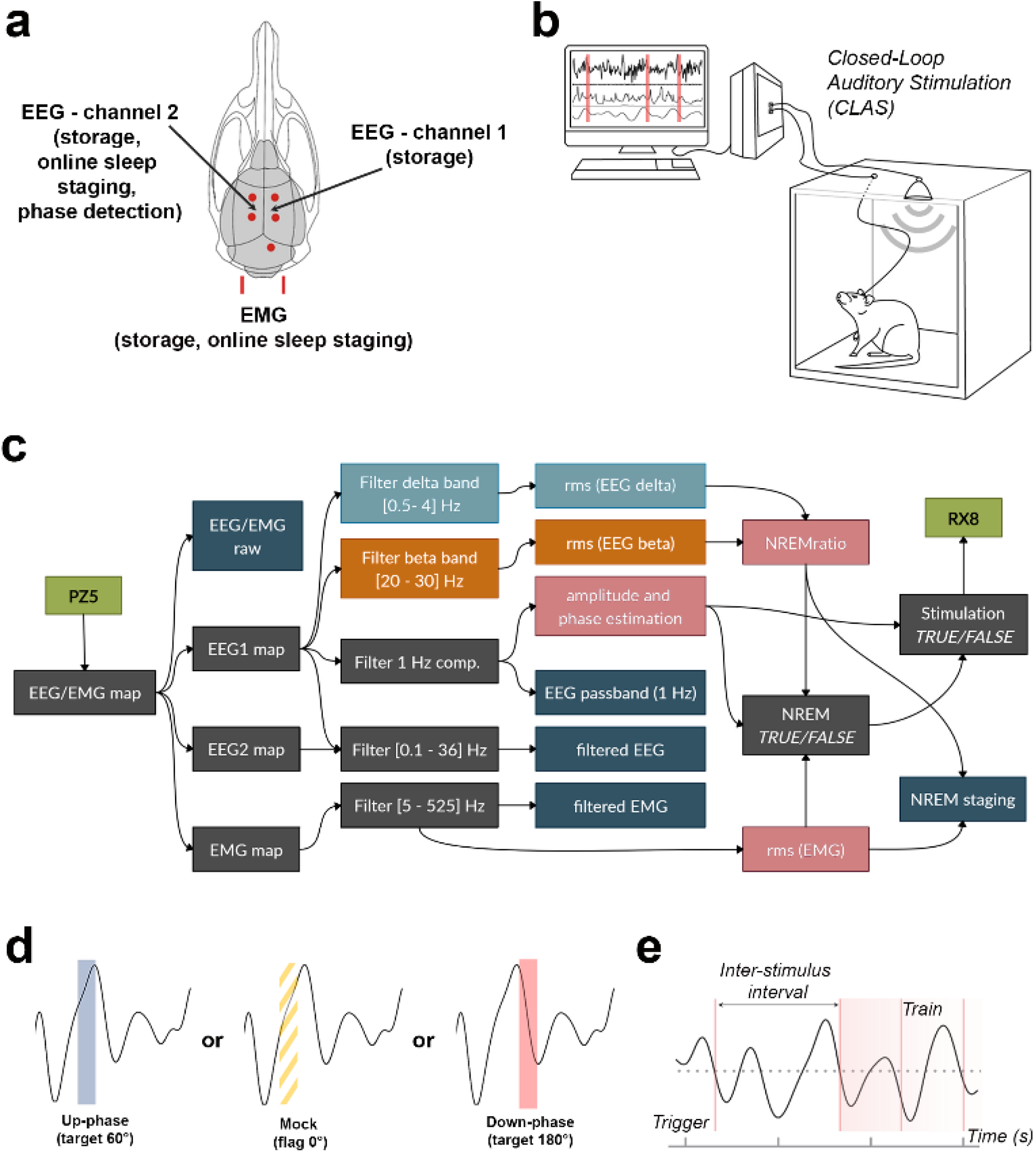
Schematic representation of EEG/EMG headpiece, CLAS concept and computational pipeline. **a.** Dorsal skull schematic of the two subdural (differential) electrodes for cortical EEG and one nuchal EMG. **b.** Closed-loop auditory stimulation setup: the system can accommodate up to 8 subjects in the same machine. **c.** Processing tree for online NREM-staging and phase-targeted auditory stimulation. EEG/EMG is sampled and amplified via PZ5 NeuroDigitizer preamplifier (TDT, USA). Left EEG (EEG1) is filtered in the delta and beta bands, and between 0.1 – 36 Hz for offline analysis; EMG signal is filtered between 5 – 525 Hz for offline analysis. Power estimations (rms(EEG delta) in orange, rms(EEG beta) in blue and rms(EMG) in pink) and amplitude of EEG 1-Hz component form the basis for detection of NREM. When NREM is identified, auditory triggers are delivered phase-locked to the EEG 1-Hz component (RX8 MULTI-I/O processor). Intermediary operations in grey and outputs for offline analysis in dark-blue. **d.** Animals were stimulated with either up-phase (60°), mock stimulation (0°, sound muted) or down-phase (180°) CLAS. **e.** Distribution of sound triggers was analysed in terms of trains of triggers (count of any size sequences of triggers 1 second or less apart) and interstimulus interval (ISI, time between triggers). EEG: electroencephalogram. EMG: electromyogram. rms: root mean square.

**Figure 2.**
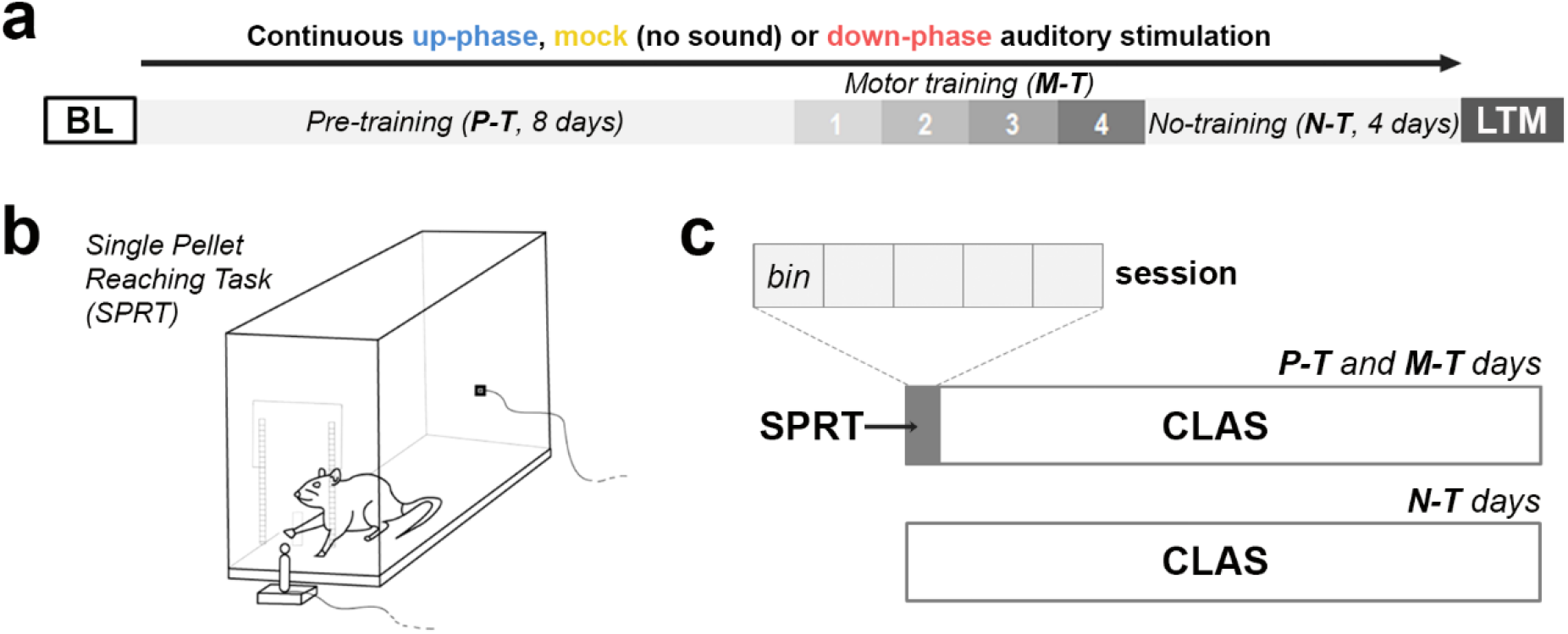
Behavioural experimental design. **a.** During the course of 16 days under continuous auditory stimulation, the animals were trained in the single-pellet reaching task. The motor-training phase started on the 9^th^ day, and continued for 4 testing days. Next, the animals moved to a no-training period, yet still under auditory stimulation, and finalized the experiment on the 16^th^ day, with a last motor assessment (LTM). **b-c.** Every day, the animals were trained in the single pellet reaching task during the first hour of the light period, and got back to the auditory stimulation chambers for the remaining time. Each training session had a cut of at 6Ominutes, divided into 5 bins of up to 12 minutes. BL-Baseline. P-T: pre-training. M-T: motor-training. N-T: no-training. LTM: long-term memory. SPRT: Single-pellet reaching task.

**Figure 3.**
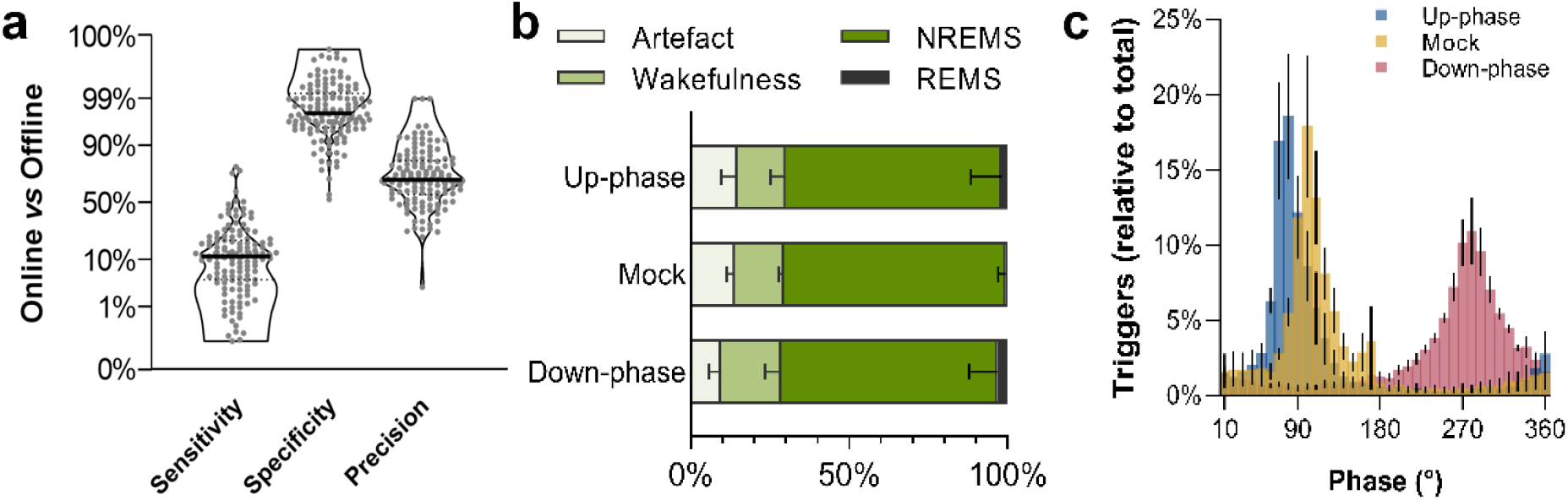
NREM sleep staging diagnostics and phase-target distribution. **a.** CLAS exhibited a 12% sensitivity. 98% specificity and 70% precision for unsupervised real-time NREM staging. **b.** Offline classification of online-labelled NREM sleep epochs per paradigm. **c.** Phase distribution at trigger onset for up-phase (targeting 65°), mock (targeting 90° but no deliver of sound) and down-phase (targeting 270°).

### CLAS modulates delta and sigma power while preserving sleep proportions over 24 hours

We analysed sleep proportions at BL, 3^rd^ day of motor training (M-T_3_), and last day of no-training period (N-T_4_). Neither up-phase nor down-phase stimulation disturbed total-NREM amounts compared to mock condition during representative days (RM-ANOVA, *time*condition* interaction, *F* (4, 32) = .789, *p* = .541, **Fig. 4a**), nor throughout the entire protocol (data not shown). We found no differences in 24-h NREM proportions within experimental conditions in M-T_3_ and N-T_4_ compared to BL (paired Wilcoxon rank tests, corrected for multiple comparisons, **Fig. 4b**). Furthermore, we observed no evidence of abnormal NREM fragmentation in any stimulation group over time (RM-ANOVA, *time*\**condition* interaction, *F* (4, 34) = 1.360, *p* = .268, **Fig. 4c**). On protocol-day M-T_3_, 24-h EEG power spectra in NREM exhibited increased delta activity in the up-phase stimulated group (**Fig. 4d**) whereas down-phase stimulated animals showed a trend towards decreased delta-frequency activity (**Fig. 4f**). Spectral density power in mock animals did not present distinctive changes on day M-T_3_ (**Fig. 4e**).

**Figure 4.**
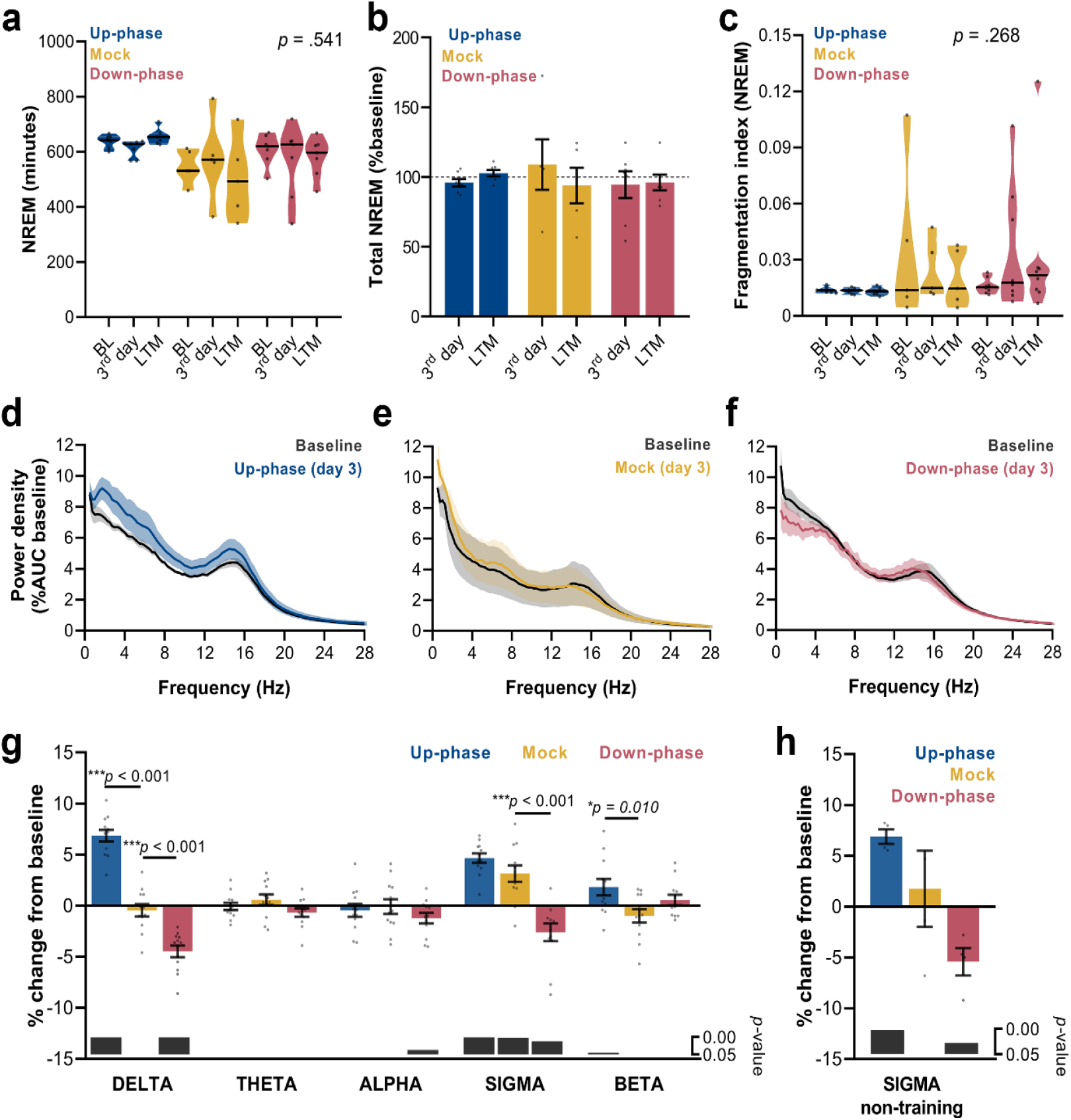
Altered delta and sigma power in stimulated animals. **a.** NREM amount is not different between groups overtime (RM-ANOVA, condition*day interaction, *F* (4, 32) = .789, *p* = .541), **b.** nor within each condition (paired Wilcoxon rank tests, corrected for multiple comparisons). **c.** Fragmentation index reveals no fragmented NREM in none of the groups overtime (RM-ANOVA, *time*condition* interaction, *F* (4, 34) = 1.360, *p* = .268). **d-f.** Average spectral power density in NREM (0.5 – 28) Hz during the 3^rd^ day of M-T, relative to baseline. **g.** 24-h spectral for the all groups as percentage change from baseline for delta (>0.5 – 4 Hz), theta (>4 – 8 Hz), alpha (>8 – 12 Hz), sigma (>11 – 16 Hz), and beta (>16 – 20 Hz) frequency bands, for all P-T and M-T days (one-sample Hotelling’s T-squared). The change in delta and beta was significantly increased during up-phase stimulation compared to mock (*\*\*\*p* < .001 and *\*p* = .010), whereas down-phase stimulation show a significant reduction in relation to non-stimulated animals both in delta and sigma (*\*\*\*p* < .001 and *\*\*\*p* < .001, respectively). Significant changes from baseline are marked as grey bars on the lower part of the chart. **h.** Sigma power in the mock group back to baseline values during N-T days (one sample t-test, *p* = .672). NREM: non-rapid eye movement. BL: baseline. AUC: area under the curve.

Notably, the stimulation design was accompanied by large inter-individual variability with respect to EEG power in NREM (**Table 1**). To highlight the global effect of each modulatory paradigm, we pooled the daily results for the various sleep-frequency bands, from P-T_1_ to M-T_3_ days, and analyzed the average change from BL. Across the spectrum, delta (*\*\*\*p* < .001), sigma (11 – 16 Hz, *\*\*\*p* < .001) and beta (16 – 30 Hz, *\*p* = .045) bands were significantly increased in up-phase stimulated subjects. On the contrary, down-phase stimulation decreased delta (*\*\*\*p* < .001), alpha (8 – 11 Hz, *\*p* = .037) and sigma (*\*p* = .012) power during P-T_1_ – M-T_3_ days, collectively. Mock animals showed altered power in the sigma band (*\*\*p* = .002, one-sample Hotelling’s T-squared, **Fig. 4g**), an apparent effect of the skill learning task performed daily. This effect was corroborated by examining sigma activity during the N-T period – 4 days without motor skill training – in all 3 groups: mock animals returned to BL indices (*p* = .672), whereas up-phase (*\*\*p* = .002) and down-phase (\**p* = .028) stimulated groups showed increased and decreased sigma activity, respectively (one sample t-test, **Fig. 4h**).

**Table 1.**
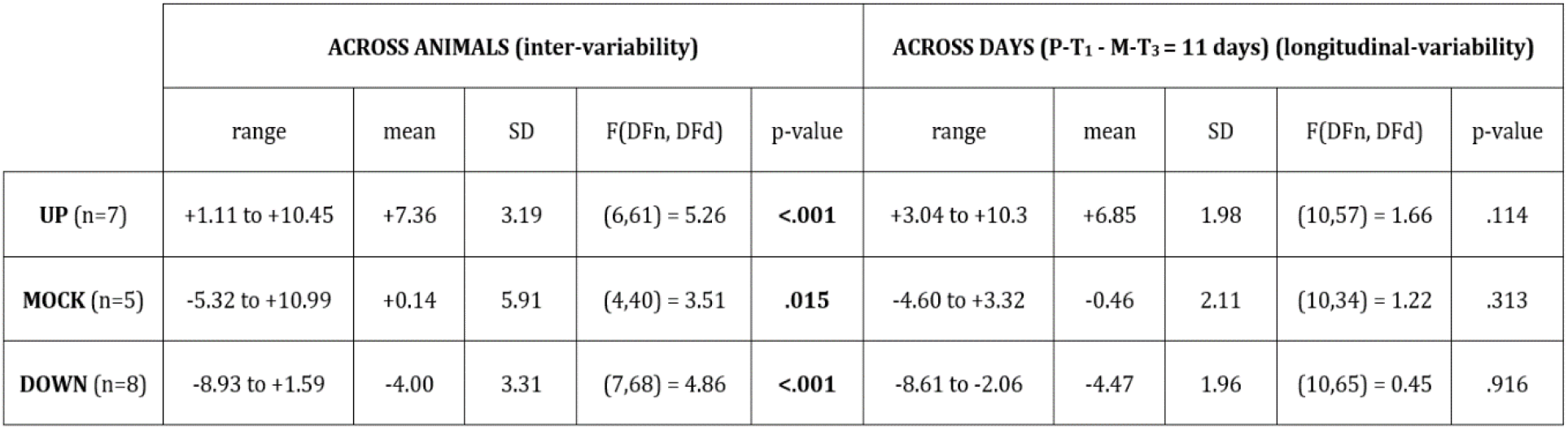
Inter-individual and longitudinal variability for delta activity (% change from BL). Differences between subjects contribute the most to the variability seen during CLAS. Values across animals represent the grand-average (P-T_1_ – M-T_3_) of delta power change from BL per animal. Values across days include a grand-average of all animals per day. P-T: pre-training. M-T: motor-training.

|  | ACROSS ANIMALS (inter-variability) |  |  |  |  | ACROSS DAYS (P-T<sub>1</sub> - M-T<sub>3</sub> = 11 days) (longitudinal-variability) |  |  |  |  |
| --- | --- | --- | --- | --- | --- | --- | --- | --- | --- | --- |
|  | range | mean | SD | F(DFn, DFd) | p-value | range | mean | SD | F(DFn, DFd) | p-value |
| **UP** (n=7) | +1.11 to +10.45 | +7.36 | 3.19 | (6,61) = 5.26 | **<.001** | +3.04 to +10.3 | +6.85 | 1.98 | (10,57) = 1.66 | .114 |
| **MOCK** (n=5) | -5.32 to +10.99 | +0.14 | 5.91 | (4,40) = 3.51 | **.015** | -4.60 to +3.32 | -0.46 | 2.11 | (10,34) = 1.22 | .313 |
| **DOWN** (n=8) | -8.93 to +1.59 | -4.00 | 3.31 | (7,68) = 4.86 | **<.001** | -8.61 to -2.06 | -4.47 | 1.96 | (10,65) = 0.45 | .916 |

### Daily time-course of delta-power changes depend on CLAS’ targeted phase, and its 24-h pattern remains stable across prolonged and continuous stimulation

To understand the daily time course of power-density in the delta-frequency band, and to appreciate the effect of chronic stimulation during NREM throughout all protocol-days, we examined the hourly change in relation to BL for each protocol-day. Up-phase stimulated animals consistently presented increased delta power throughout the day, with larger increases during the light phase (up to ~ 23% in some days), an effect that persisted until the end of the protocol (**Fig. 5a**). Mock animals oscillated around zero-change value (**Fig. 5b**), in concordance with the notion of a reorganized sleep-wake pattern, due to the 1-h long SPRT task and food-entrainment activity (Northeast et al., 2019), the later notably present in the other groups as well. Down-phase stimulated animals exhibited reduced delta power for the majority of the time, with greater hourly decreases during the dark period, down to ~ −17% (**Fig. 5c**). We evaluated the time course of delta power from P-T_8_ to M-T_1-4_ days to explore the effect of stimulation before and after the SPRT hour. In up-phase stimulated animals, the time-course analysis of delta activity revealed significantly higher power during the hours immediately after the testing session compared to BL (multiple paired t-tests, corrected with Holm-Sidak method for multiple comparisons; **Fig. 5d**). This finding overlapped with the animals’ largest daily drop in homeostatic sleep pressure (*see BL curve*, **Fig. 5d**). While mock animals did not show differences during the aforementioned stimulation days compared to BL (**Fig. 5e**), down-phase stimulated animals presented the largest decrease compared to BL during the hours immediately before the SPRT (multiple paired t-tests, corrected with Holm-Sidak method for multiple comparisons; **Fig. 5f**). Contrast between delta activity during the single-pellet reaching task session’s preceding and succeeding hours for up- and down-phase stimulation groups shows significantly modulation levels for down-phase stimulated animals in the end of the dark period, and for up-phase stimulated animals immediately after the testing session, during the light period (**Fig. 5g**).

**Figure 5.**
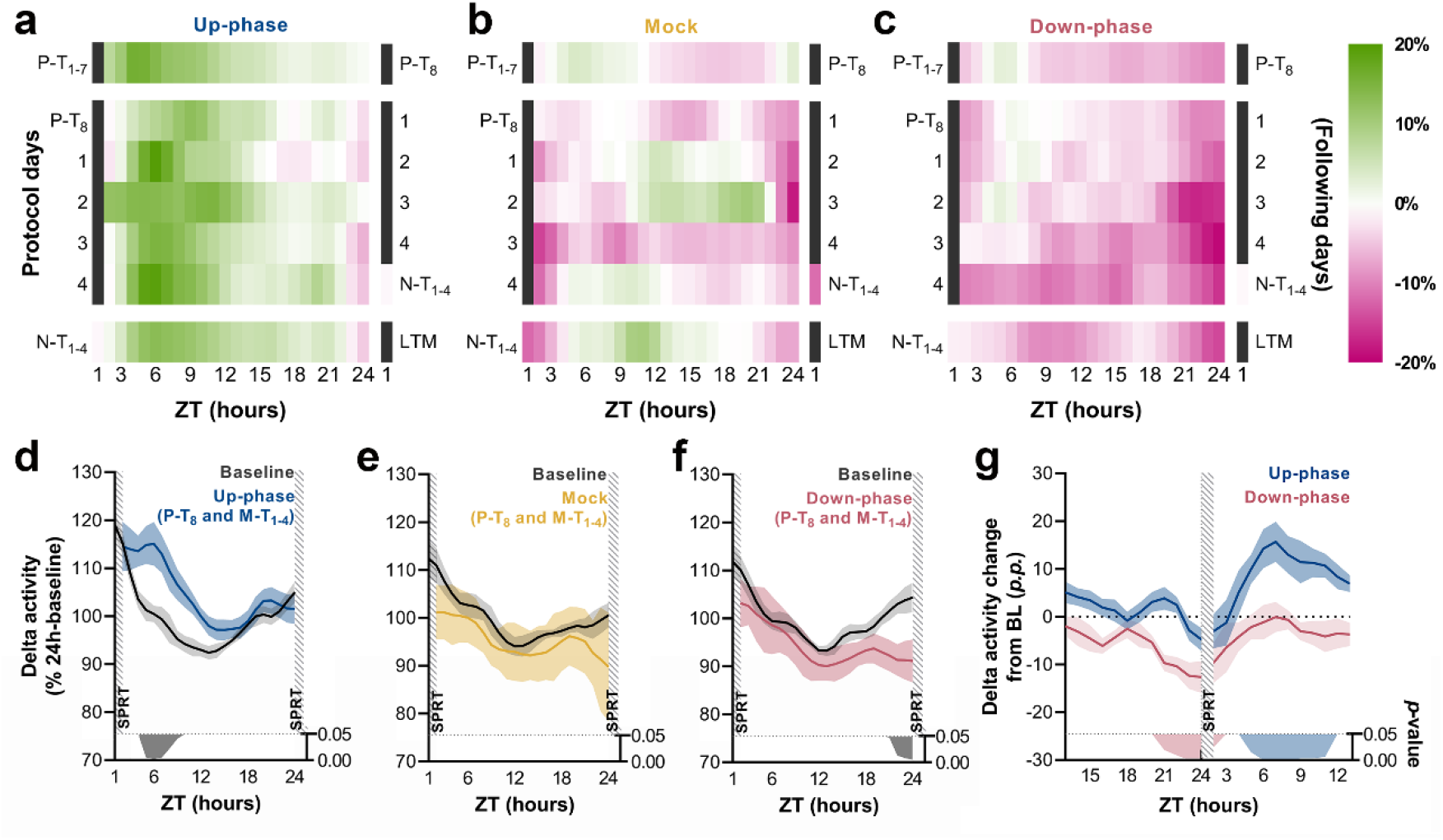
Time-course of delta activity changes from baseline during all protocol days and motor-training grand-average. **a-c.** Delta activity (0.5 – 4 Hz) changes from baseline, in 1-h bins, for up-phase, mock, and down-phase stimulated animals. P-T and N-T days were condensed into grand averages (top and bottom rows, respectively). Individual M-T days are shown in the middle rows. Hours in black represent the single-pellet reaching task training sessions, numbered with the respective protocol-day. On the right side of each heat-map, the training sessions of the following protocol-day are enumerated. **d-f.** Time-course of hourly delta-activity during BL and motor-training days, collectively (coloured line represents the grand-average of training sessions P-T_8_ to M-T_4_). **g.** Contrast between delta activity during the single-pellet reaching task session’s preceding and succeeding hours for up- and down-phase stimulation groups, plotted in percentage points from BL. Grey/coloured shadows bellow plots D-G represent significant time-points (multiple t-tests, Holm-Sidak correction for multiple comparisons). ZT: zeitgeber. P-T: pre-training. M-T: motor-training. N-T: no-training. LTM: long-term memory. SPRT: single-pellet reaching task. p.p. percentage points.

### CLAS modulates sigma power in a targeted-phase dependent manner

CLAS targeting slow waves also altered sigma power (**Fig. 4g**). The hourly time course of sigma-power changes showed a steady boost throughout the experiment, up to ~ 12% (**Fig. 5-figure supplement 1a**). Mock animals exhibited non-significant power changes (range: ~ −10% to ~ 14%) across all stimulation days (**Fig. 5-figure supplement 1b**). The decrease in sigma power in the down-phase stimulated animals reached depressions down to ~ −12% during the light period, immediately after the test (**Fig. 5-figure supplement 1c**). When we investigated the combined effect of CLAS on P-T_8_ and M-T_1-4_ days immediately after the training sessions, we found no significant change in sigma power in the up-phase stimulated and mock groups compared to their BL (multiple paired t-tests, Holm-Sidak method for multiple comparisons, **Fig. 5-figure supplement 1d-e**). On the contrary, down-phase stimulation reduced sigma power compared to BL, particularly during the hours following the motor training task (multiple paired t-tests, Holm-Sidak method for multiple comparisons, **Fig. 5-figure supplement 1f** and **Fig. 5-figure supplement 1g**).

### Up-phase CLAS preserves the natural proportion of same-size trains of triggers, whereas down-phase CLAS prolongs the interval between triggers

Next, we examined the frequency of trains of triggers (interstimulus interval (or ISI) ≥ 0.8 s): mock animals show a skewed distribution, with single triggers representing ~ 65% of all sequences of triggers and ~ 43% of all triggers in NREM (*M* = 2384, *SD* = 2296, per 24 h). Up-phase stimulated animals showed no difference in proportion of trains of any size when compared to the mock group (**Fig. 6a**), however they overall received more triggers per day (*M* = 5862, *SD* = 2214). Conversely, down-phase stimulated subjects presented a different pattern of stimulation (2-way ANOVA, *condition*train-size* interaction, *F* (8, 80) = 6.72, \*\*\**p* < .001, **Fig. 6a**), with significant higher rate of single triggers (\*\*\**p* < .001, ~ 58% of all triggers in NREM per 24 h, *M* = 2462, *SD* = 1686) and less trains of 2 triggers (*\*p* = .020), hinting on the disruptive effects of down-phase targeting on slow waves. Corroborating these results, the analysis of ISIs (2-way ANOVA, *condition*ISI* interaction, *F* (16, 144) = 1.80, *\*p* = .037) showed that up-phase targeting preserved the proportion of same-length non-stimulated periods seen in the analysis of muted-sound triggers in non-stimulated subjects, whereas down-phase stimulation provided a distribution of triggers containing significantly less density of ISIs < 2 s (**Fig. 6b**).

**Figure 6.**
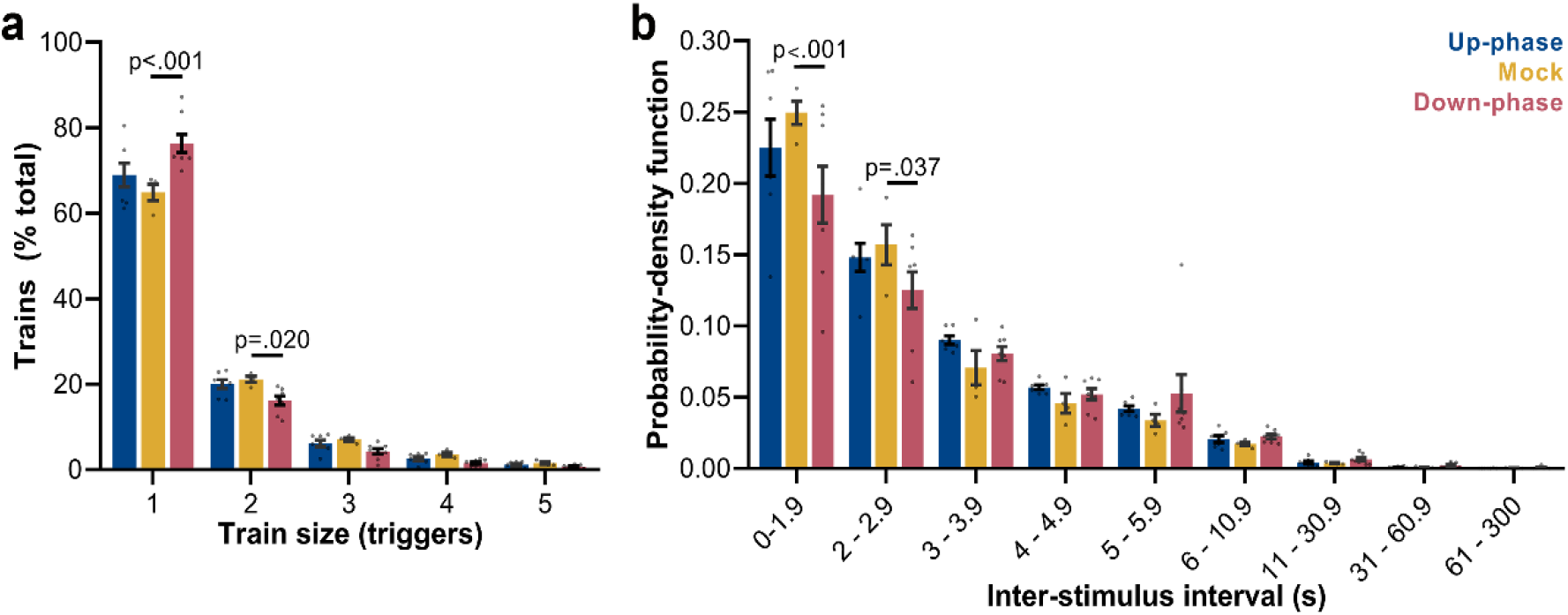
Trigger’s entrainment and distribution. **a.** Histogram for the train of pulses, up to 5 consecutive stimuli, spaced by no more than 1s (2-way ANOVA, *condition*train size* interaction, *F* (8, 80) = 6.72, *\*\*\*p* < .001). **b.** Histogram for the time between triggers, organized in a probability density function for different bin sizes (2-way ANOVA, *condition*interstimulus interval* interaction, *F* (16, 144) = 1.80, *\*p* = .037).

### Up-phase CLAS mildly entrains EEG slow-waves whereas stimulation during the down-phase disturbs it

Analysis of event-related potentials (ERP, filtered in the 0.5 – 2 Hz frequency band), i.e. all waveforms receiving triggers during NREM, revealed a small slow-wave cycle following the endogenous cycle that triggered the stimulation in up-phase CLAS (**Figure 6-figure supplement 1a**), while down-phase targeting appeared to completely disrupt the slow-wave negative half-cycle (**Figure 6-figure supplement 1b**). This is evidenced by the amplitude contrast between the two conditions for two succeeding slow-wave troughs (**Figure 6-figure supplement 1c**). Although not statistically significant, we observed trends for lower amplitude for both the depolarized up-state (unpaired t-test, *p* = .065) and the hyperpolarized down-state (unpaired t-test, *p* = .058) in down-phase stimulated animals compared to up-phase stimulated rats.

### Phase-targeted CLAS modulates success rate of motor learning in the SPRT

We first analyzed the raw counts for attempts, successes, failures and drop-ins. Attempts’ counts had high dispersion for every M-T day, and although they showed considerable qualitative clustering at day 4 (**Fig. 7a**), their high dispersion did not allow differentiation of the groups. The lack of apparent interaction between *group* and *day* was confirmed by the negative-binomial distribution model (*Х^2^* (2, *N*=21) = 0.25, *p* = .883). Increase in the incidence rate ratios (IRR) of attempts across M-T days did not achieve significance for mock animals (*n* = 6; *IRR* = 1.07; *p* = .126), but revealed a tendency of growth in the up-phase stimulated group (*n* = 7; *IRR* = 1.08; *†p* = .061) and a significant increase in the down-phase stimulated group (*n* = 8; *IRR* = 1.10; *\*p* = .014). Successes’ count grew in all groups (*Х^2^* (1, *N*=21) = 50.08, *\*\*\*p* < .001), but without significant interaction (*Х^2^* (2, *N*=21) = 0.55; *p* = .761, **Fig. 7b**), rendering neither up- (*p* = .613) nor down-phase (*p* = .123) stimulated groups significant differences from mock animals (*n* = 6). Failed attempts’ counts showed differences between groups dependent on day (*Х^2^* (2, *N*=21) = 8.29, *\*p* = .016). This result reflects the decrease in the incidence of fails in the up-phase stimulated group (*IRR* = 0.75; *\*\*\*p* < .001), but not in the down-phase group (*IRR* = 1.01, *p* = .864), with mock intermediate to both (*n* = 6; *IRR* = 0.85, *†p* = .054, **Fig. 7c**). A collective reduction in drop-in counts (the pellet is grasped but dropped during arm retraction) is found over time (*Х^2^* (1, *N*=21) = 4.42, *\*p* = .035), without differences between groups (data not shown). In summary: a discernible improvement in the up-phase stimulated group is evidenced by a significant decrease in failed attempts and a minor increase in successes over time, whereas down-phase stimulated animals attempt significantly more as protocol progresses, but are not as successful, and display a high failure count relative to total attempts. Mock animals (*n* = 6) sit somewhat in between the learning curves of stimulated groups, marked by a moderate increase in successes and a tendency to fail less over the 4 M-T days. The evaluation of success rate (SR), the number of successes per total attempts, demonstrated that down-phase stimulated subjects present a significantly worse learning progression when contrasted with mock subjects (*n* = 6, *Х^2^* (2, *N*=21) = 17.50, *\*\*\*p* < .001; Tukey post-hoc comparison *up*mock: *p* = .040, **Fig. 7d**). SR at LTM-day did not significantly differ from M-T_4_ in any of the groups (2-way ANOVA, *time*condition, F* (2, 17) = 0.12, *p* = .887).

**Figure 7.**
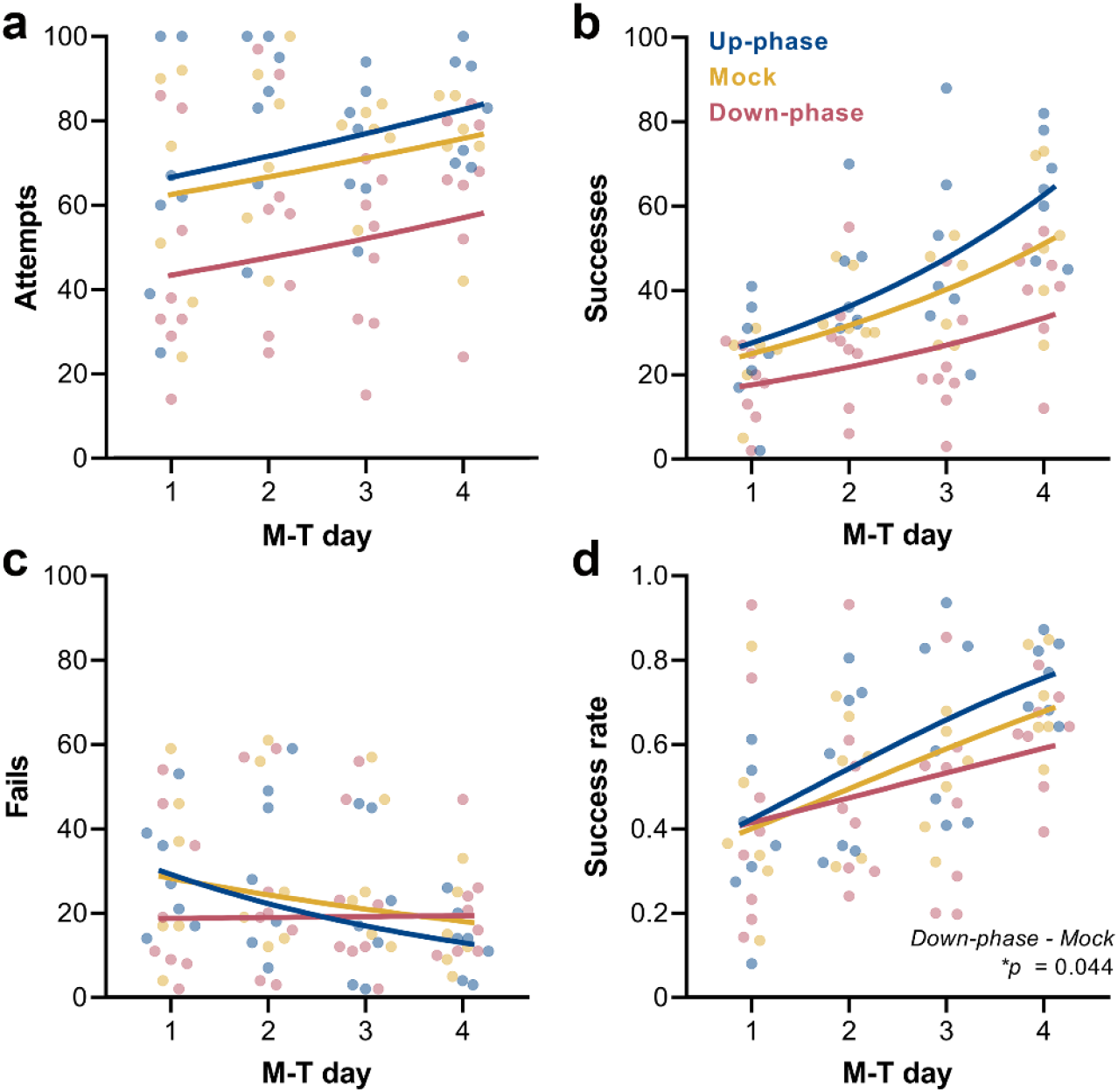
Up-phase stimulated animals attempt more, succeed more and fail less over time, whereas down-phase stimulated rats show a significantly reduced success-rate. **a.** Attempts’ counts showed high dispersion for every M-T day. **b.** Successes’ grow significantly in all groups (*Х^2^* (1, *N*=21) = 50.08, *\*\*\*p* < .001), but no differences were found between the stimulated groups and mock. **c.** Groups showed significant differences in fail counts over time (*Х^2^* (2, *N*=21) = 8.29, *\*p* = .016). Up-phase stimulated group saw a significant reduction in fail counts (*IRR* = 0.75, *\*\*\*p* < .001), but down-phase group did not (*IRR* = 1.01, *p* = .864). **d.** By observing SR (number of successes in total attempts), down-phase stimulated subjects show a significantly worse learning progression when contrasted with mock subjects (*Х^2^* (2, *N*=21) = 17.50, *\*\*\*p* < .001; Tukey post-hoc comparison *up*mock: *p* = .040). M-T: motor-training.

### Up-phase stimulated animals show daily intra-session improvements, while down-phase stimulated animals attempt less towards the end of the session

Compartmentalizing each training session into 12-min bins (or 20 pellets, whichever comes first) revealed that up-phase stimulated and mock subjects maintained the same rate of attempted grasps across bins (RM-ANOVA, *condition*bin* interaction, *F*(2, 18) = 4.03, *\*p* = .036, **Fig. 8a**), while down-phase stimulated animals attempted significantly less towards the end of the session (*first-bin*last-bin, *p* = .022) and significantly less than control animals during the last bin (*down*mock, *p* = .037). When relativizing number of attempts per bin by the total number of attempts within the session (RM-ANOVA, *condition*bin* interaction, *F* (2, 18) = 5.24, *\*p* = .016, **Fig. 8b**), down-phase stimulated animals again show a sharp decrease towards the end of the session (*first-bin*last-bin, **p* = .004). Attempting speed (number of attempts per minute) also showed a decline from the 1^st^ to the 5^th^ bin in the down-phase stimulated subjects, but failed to reach significance (RM-ANOVA, *condition*bin* interaction, *F* (2, 18) = 1.33, *p* = .291, **Fig. 8c**). Although the intra-session attempt rate in the up-phase stimulated group remained constant from the beginning to end of training sessions, there was an increase in successful attempts from the 1^st^ to the 5^th^ bin in this group in all M-T days. Conversely, down-phase stimulated animals were always slightly less successful towards the end of each session (**Fig. 8d**), although this observation might be explained by the reduction in attempts along the session duration.

**Figure 8.**
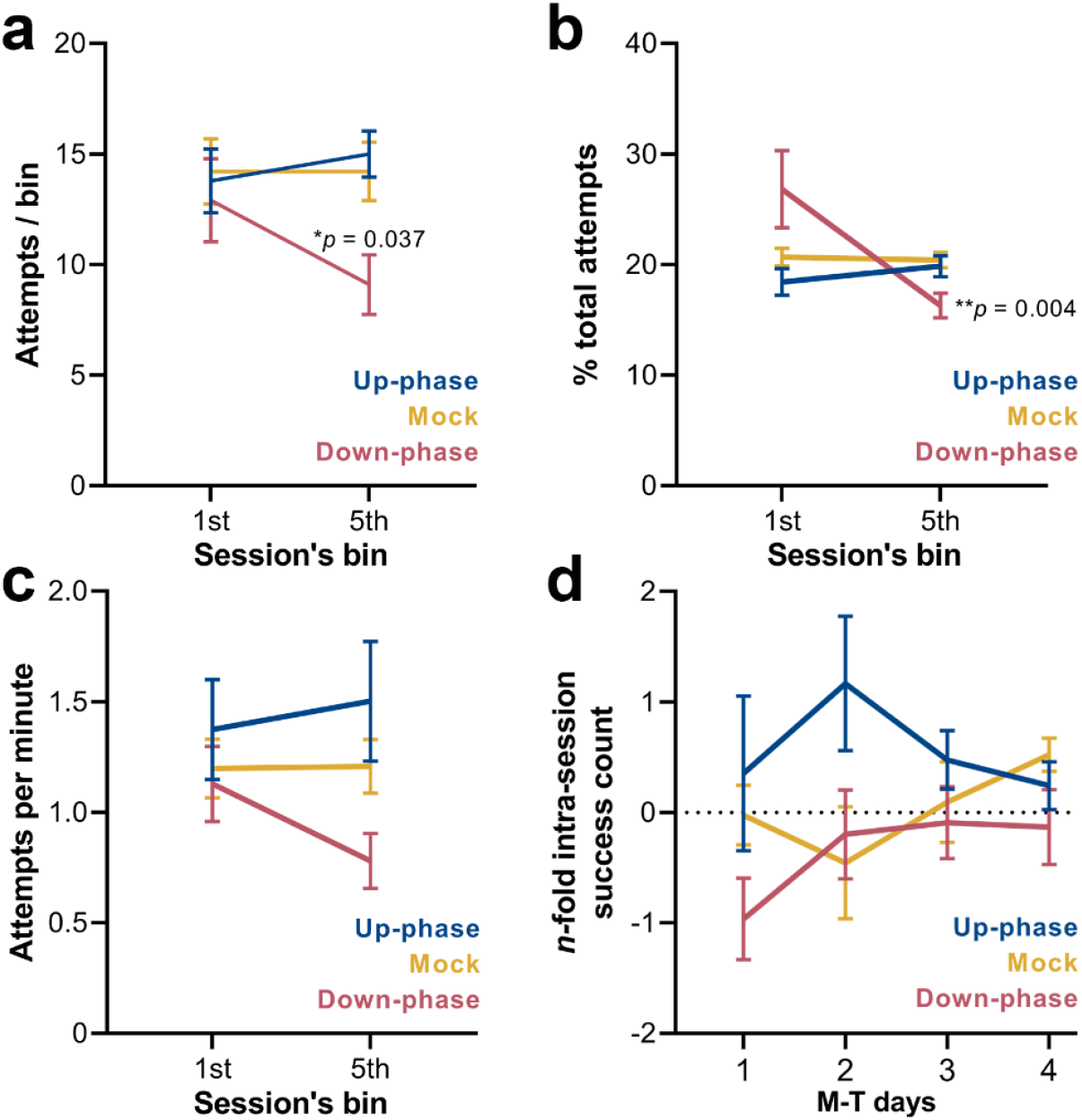
Down-phase stimulated animals show a reduction in the engagement with the task towards the end of the session. **a.** Down-phase stimulated animals attempt significantly less than control animals during the last bin (RM-ANOVA, *condition*bin* interaction, *F* (2, 18) = 4.03, *\*p* = .036; *down*mock, *p* = .037). **b.** In relation to the proportion of attempts per bin to the total number of attempts, down-phase stimulated animals again show a sharp decrease towards the end of the session (RM-ANOVA, *condition*bin* interaction, *F* (2, 18) = 5.24*, *p* = .016; *first-bin*last-bin, **p* = .004). **c.** Attempting speed shows a non-significant decline from the 1_st_ to the 5_th_ bin in the down-phase stimulated subjects (RM-ANOVA, *condition*bin* interaction, *F* (2, 18) = 1.33, *p* = .291). **d.** Up-phase stimulated animals show a consistent increase in successful attempts from the 1_st_ to the 5_th_ bin, in all M-T days. Conversely, down-phase stimulated animals were always slightly less successful towards the end of each session, although this might be due to the drop in attempts within the last testing window (plotted as the log2 of the fold-change of the number of successes of the last bin to the first bin of a session: −1 represents a 50% drop in successes, while 1 represents twice more successes from first bin). M-T: motor-training.

### *Number of triggers correlate with magnitude of delta-power changes, and both delta and sigma changes positively correlate with success-rate at the end of the* M-T *period*

To investigate whether the number of sound triggers influences the extent of delta-power modulation, we extracted the number of triggers over each 24-h period during the M-T phase for all stimulated animals. The magnitude of changes in the delta-frequency band was moderately correlated with the number of delivered stimuli, both for up- (Spearman’s *rho* = 0.43, *\*p* = .022, **Fig. 9a**) and down-phase stimulated animals (Spearman’s *rho* = −0.41, *\*p* = .026, **Fig. 9a**). We also found a significant positive association between 24-h changes in delta and sigma power (Spearman’s *rho* = 0.57, \**\*p* = .004, **Fig. 9b**), reflecting the impact of CLAS on the two most dominant features during NREM sleep. Subsequent analysis between 24-h average delta change (all training-phase days condensed) and SR at the end of the training phase revealed a moderate positive association between modulation of delta activity and SPRT performance (Spearman’s *rho* = 0.41, *\*p* = .034, **Fig. 9c**). Likewise, sigma-activity changes positively correlate with SR scores (Spearman’s *rho* = 0.53, *\*p* = .007, **Fig. 9d**).

**Figure 9.**
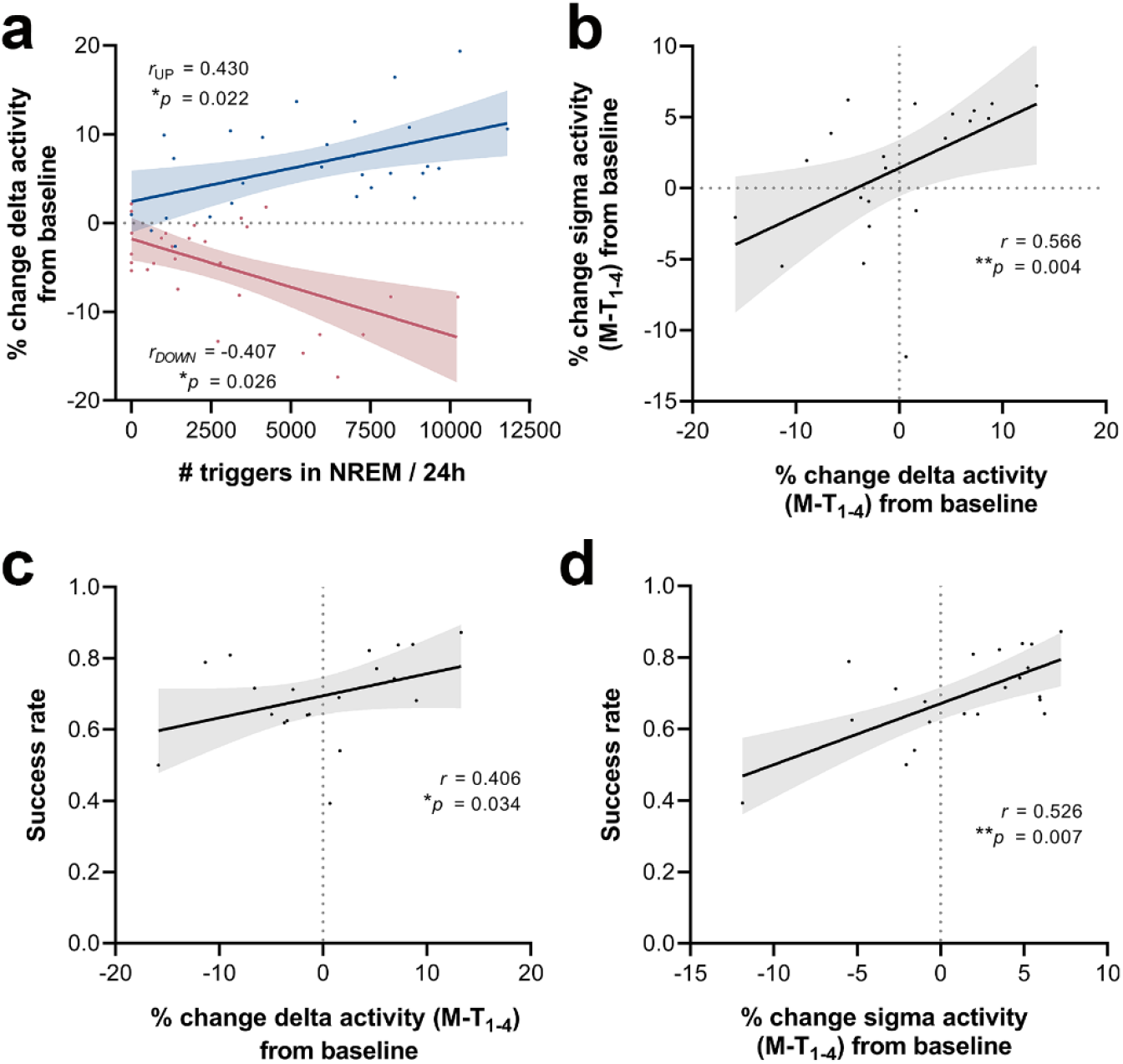
Correlations between number of triggers, delta-activity change from baseline, sigma-activity change from baseline and success-rate. **a.** The magnitude of changes in the delta-frequency band was moderately correlated with the number of stimuli delivered, both for up- (Spearman’s *rho* = 0.43, *\*p* = .022) and down-phase stimulated animals (Spearman’s *rho* = −0.41, *\*p* = .026). **b.** In turn, 24-h grand-average (M-T days) of delta-activity changes from BL correlated significantly with sigma-activity changes in the same period (Spearman’s *rho* = 0.57, *\*\*p* = .004). **c-d.** Moderate positive associations between the SR scores achieved by all subjects in the last motor assessment (M-T_4_) and the global changes in delta-activity (Spearman’s *rho* = 0.41, *\*p* = .034) and sigma-activity (Spearman’s *rho* = 0.53, *\*p* = .007), from all days preceding the sessions of the motortraining phase, collectively (P-T_8_, M-T_1_, M-T_2_ and M-T_3_). M-T: motor-training.

## Discussion

### Conceptual and technical validation of rodent CLAS: Chronic continuous phase-targeted CLAS specifically and steadily modulated delta and sigma power

This is the first report of successful phase-targeted CLAS in rodents. With this paradigm, we delivered auditory triggers to healthy rats continuously during a period of 16 days. We observed a mild but specific mean increase of 6.9% (up to ~ 23% during the light period) in delta power per 24 h on stimulation days compared to BL. This effect was stable over the entire stimulation period evaluated, with no signs of decay nor facilitation effects over time. Reverse effects (*M* = - 4.5%, down to −17.5% by the end of the dark period) were observed with down-phase stimulation.

Notably, application of CLAS in human subjects has often failed to demonstrate an overall inter-night effect (unperturbed night *vs* stimulated night) of stimulation on spectral power (Papalambros et al., 2017; Papalambros et al., 2019), while comparison between ON and OFF stimulation windows within the same night yielded significant stimulation effects that positively correlated with subsequent memory performance (Ong et al., 2016; Papalambros et al., 2017). In this study, we provide first evidence of a long-term effect when comparing unperturbed 24 h (BL) *vs* stimulated 24 h in a paradigm of continuous stimulation, somewhat challenging the proposition that the effect of CLAS is mediated by qualitative transient reorganization of SWA rather than of a general SWS or SWA enhancement (Papalambros et al., 2017). CLAS elicited, however, a modest effect in both up- and down-conditions in our rats, without evidence of either build-up or adaptation mechanisms to regulate the impact of the stimulation across the days. Interestingly, chronic sleep deprivation studies in rodents demonstrate a comparable effect, suggesting that strong homeostatic compensatory mechanisms may be at play (Kim, Hwang, Strecker, Choi, & Kim, 2020; Leemburg et al., 2010).

To achieve a greater effect, further refined CLAS paradigms must be evaluated. For instance, although the CLAS protocol employed in this study accomplished 70% NREM precision and clear phase-target accuracy, future NREM online-staging and CLAS pipelines shall be optimized in terms of detection of particular EEG events, such as slow oscillations, spindles or arousals. Additionally, triggers would ideally be delivered only during sustained NREM sleep, but motion behaviours with a rhythmic nature (e.g. grooming, feeding, and drinking) may have been major factors contributing to wakefulness and artifact epochs being identified online as NREM sleep, and consequently reducing precision. Thus, further artifact-rejection features shall be considered, such as auxiliary electromyogram (EMG) information (e.g. piezo-electric sensors), especially in long-term experiments where attrition of implanted EEG/EMG headsets is unavoidable. Also, very strict thresholds for NREM sleep online identification drastically limited sensitivity of the algorithm (12%). Moreover, implementation of phase-locked loop (PLL) algorithms in human studies, has proven to offer superior results and shall be attempted in rodent CLAS in the future (Ferster et al., 2019; Santostasi et al., 2016). Nonetheless, the CLAS paradigm introduced here produces stable sleep modulation effects over time and successfully replicates the timing accuracy so far obtained in some methods of human CLAS (Fattinger et al., 2019; Papalambros et al., 2017). Also like in humans studies (Henin et al., 2019; Krugliakova, Volk, Jaramillo, Sousouri, & Huber, 2020; Papalambros et al., 2017), we observed an additional modulatory effect of CLAS on power in the sigma-frequency band in rats. Sigma modulation was, just like CLAS-dependent delta changes, phase-target specific and steady over the 16 days of stimulation.

We explored the size of trains of triggers and interstimulus intervals (ISIs) (Santostasi et al., 2016) — proxies for length and distribution of delta wave trains — to explore whether CLAS altered natural slow wave patterns and if this could be the potential underlying process by which CLAS exerted its modulatory effect over delta power. In comparison to mock, down-phase stimulated animals showed an increased number of single-triggers, decreased proportion of trains of 2 triggers, and decreased density of <2 s ISIs, in contrast with maintained levels in the up-phase stimulated group, potentially indicating that down-phase CLAS has disarrayed the natural pattern of delta wave trains.

In agreement with the apparent preservation of the slow wave pattern with up-phase CLAS, and opposite effect with down-phase CLAS, analysis of event-related potentials (ERPs) indicated the occurrence of a post-trigger slow wave when targeting 60°, whereas a post-trigger pattern disruption arises from 180° target stimulation. Peak-to-peak amplitude did not differ between up- and down-phase groups, although we cannot reject a possible tapering of the physiological response from stimuli outside of the desired phase. Taken together, these findings suggest that CLAS of rodent slow waves may be altering the natural slow waves’ pattern rather than significantly affecting their amplitude.

### Functional validation rodent CLAS: Altered behavioral performance upon phase-targeted CLAS in healthy rats

Phase-targeted CLAS produced significantly altered learning performance in the SPRT in healthy rats. Moreover, we found that the mean delta activity change in motor-training days 1–4 in relation to BL correlated positively with success rate on motor-training day 4 in the SPRT. Through the positive correlation found between the number of triggers — i.e. extent of closed-loop auditory stimuli delivered — and delta-power changes in up- and down-phase groups, we confirmed indeed that delta-activity changes during motor-training days 1–4 are actually driven by CLAS. Overall, these results suggest that CLAS-dependent delta-activity levels drive next-day performance in healthy rats. Furthermore, we observed that the level of proficiency achieved at motor-training day 4 was maintained in all groups after 5 days without training. This observation suggests that CLAS modulated sleep-dependent skill acquisition and retention only immediately following a motor-training session. In short, without prior active task engagement or performance saturation (Kvint et al., 2011), up-phase CLAS *per se* seems insufficient to induce a significant performance enhancement, whereas there was no evidence of skill decay induced by down-phase CLAS either. Two sleep-dependent mechanisms may explain the observed effect of modulated slow waves on behavioral performance in our paradigm. Sleep-mediated synaptic downscaling, or recovery, after learning has been deemed critical to maintaining behavioral performance (Fattinger et al., 2017). Thus, it is conceivable that targeting slow waves with CLAS might have altered (up-: increased, down-: decreased) the normal recovery process after learning during wakefulness in our rats. In support of this argument, we observed that the number of attempts in the down-phase group was lower than in the other groups at motor-training day 1, indicating that before any memory consolidation process took place, these animals already performed worse than their counterparts did. On the other hand, memory consolidation, a process at least partially mediated by spindles (Diekelmann & Born, 2010) in which specific synapses are strengthened, is also potentially sensitive to CLAS (Krugliakova et al., 2020; H.-Viet V. Ngo et al., 2013; Papalambros et al., 2017). In fact, we did observe positive correlations between sigma — potentially a proxy for spindles (Holz et al., 2012) — during motor-training days 1-4 and success rate on motor-training day 4. The two theories, however, are certainly not mutually exclusive (Klinzing, Niethard, & Born, 2019; Wilckens, Ferrarelli, Walker, & Buysse, 2018) and sleep-dependent memory benefit could well complementarily rely on them both. Our tool, thus, may be utilized to disentangle the exact effects of CLAS-modulated SWS on rodent cognition. Further, CLAS can be strategically combined with other tools, such as multiunit-activity recording, and provide crucial insight into the underlying neuronal mechanisms of auditory modulated slow waves and sigma (Krugliakova et al., 2020), and their functional role on learning and memory.

### Limitations

An experimental limitation is the fact that we did not actively apply randomization, nor exclusion criteria, in relation to the paw-preference in our experimental groups of all-male rats. Thus, as we always used the left hemisphere EEG derivation for CLAS, distinct local network dynamics determined by paw dexterity (Hanlon, Faraguna, Vyazovskiy, Tononi, & Cirelli, 2009; Vyazovskiy & Tobler, 2008) may have potentially affected closed-loop stimulation features in an uncontrolled manner in left- or right-pawed experimental subjects. Larger group numbers allowing for exclusion or efficient randomization of this confounder variable shall be considered in future efforts. Lastly, in our temporal analysis of the delta-power changes per 24 h, we observed a clear pattern showing that animals in all groups present distinctively decreased delta power levels in the few hours before the behavioral test. Whether this activity pattern, likely attributable to food anticipation (Northeast et al., 2019), might have compounded with our CLAS-specific delta power findings remains to be determined and, ideally, further controlled in future experiments. Finally, in relation to the lack of effect of phase-targeted rodent CLAS on ERP amplitudes, as others demonstrated earlier, the broad range achieved by our phase-tracking algorithm might have contributed to the observed variable effects of CLAS across animals at the ERP level, ultimately leading to high inter-individual variability on delta activity modulation, result that agrees with previous studies using non-adaptive prediction algorithms (Cox et al., 2014; Fattinger et al., 2017; Fattinger et al., 2019; H.-Viet V. Ngo et al., 2013). Employing adaptive feedback methods based on phase-locked loops (at times associated with amplitude thresholds), producing optimal phase-prediction and inducing more specific and systematic effects on SWS (Ong et al., 2016; Papalambros et al., 2017; Papalambros et al., 2019; Santostasi et al., 2016), shall be assessed in rodents next.

Altogether, our report presents the first successful implementation of phase-targeted CLAS in rats, demonstrating that phase-targeted continuous chronic CLAS is feasible, specific, and effective in rodents. Overall, this novel tool provides successful grounds to not only further study the substrate mechanisms by which CLAS affects slow wave activity and associated outcomes, such as cognition, but to additionally assess its effect on models of brains disease. Ultimately, CLAS of slow waves in rodents may help pave the way to the clinical implementation of this novel technique in the context of brain/body disorders that may be targeted with sleep-based therapeutics (Arora & Taheri, 2015; Cook, Ferry, & Tran, 2020; Fung et al., 2011).

## Materials and methods

### Animals, surgeries and husbandry

To develop this tool, we used 23 young-adult male Sprague-Dawley rats (Charles River, Italy) weighing 200-220g and group-housed them in standard IVC cages (T2000) prior to interventions. In all animals, we surgically implanted electrodes for continuous recording of electroencephalography and electromyography (EEG/EMG) as described previously(Büchele et al., 2016). Briefly, we inserted four stainless steel miniature screws (Hasler, Switzerland), one pair for each hemisphere, bilaterally into the rats’ skull following specific stereotactic coordinates: the anterior electrodes were implanted 3mm posterior to bregma and 2mm lateral to the midline, and the posterior electrodes 6mm posterior to bregma and 2mm lateral to the midline. For monitoring of muscle tone, we inserted into the rats’ neck muscle a pair of gold wires that served as EMG electrodes (**Fig. 1a**). All electrodes were connected to stainless steel wires, further connected to a headpiece (Farnell, #M80-8540842, Switzerland) and fixed to the skull with dental cement. We performed all surgical procedures under deep anesthesia by inhalation of isoflurane (4.5% for induction, 2.5% for maintenance), and subsequent analgesia with buprenorphine (s.c., 0.05 mg/kg). Following surgery, we housed the animals individually for a minimum of 14 days for recovery, with food and water available ad libitum, and handled them daily for postoperative monitoring, body weight checkup and familiarization with the experimenter. The animal-room temperature was maintained at 22 – 23 °C, and animals were kept on a 12h light-dark cycle. 2 weeks after surgery, we transferred the animals, still individually, to custom-made acrylic-glass cages (26.5 × 42.5 × 43.5 cm). Each cage was positioned inside a sound-attenuated chamber, built in house, for EEG/EMG recording and auditory stimulation (**Fig. 1b**). All procedures were approved by the veterinary office of the Canton Zurich (license ZH231/2015) and conducted in accordance with national and institutional regulations for care and use of laboratory animals.

### Sound insulation chambers

Eight 40 × 50 × 60 cm sound insulated chambers were design and built in-house with the purpose to simultaneously record physiological signals and deliver auditory stimulation to eight individual subjects at a time. Each chamber (**Fig. 1-figure supplement 1a**) was built using the following components:

- PVC boards (1 cm thickness, 0.60 g/cm^3^; KÖMATEX_®_, Röhm, Switzerland), to form a lightweight but robust structure.
- One layer of absorptive closed-cell foam (2cm thickness, 0.02 g/cm^3^, PU WAVE TOP, Swilo, Switzerland), covering the entire inner surface of the chamber, and a soundproofing closed-cell foam with protective skin and heavy rubber layer (3,2 cm thickness, 0.03 g/cm^3^, PU SKIN, Swilo, Switzerland) on the outer surface for reduction of sound and vibration transmissions.
- 1 fully demountable door, with the same isolation as described above.
- LED strips (Philips LightStrips Flex Color), attached around the chamber’s ceiling, were connected to an intensity controller and dedicated timer for light – dark cycle. The light was evenly diffused using a translucent acrylic plate (3mm thickness, *T*(D_65_) = 72%, Plexiglas_®_, Röhm, Switzerland).
- An additional infrared LED was also mounted to allow for continuous infrared video recordings.
- Passive slip-ring (Dragonfly Research, USA) for EEG/EMG recordings in freely moving animals.
- Two low-noise fans (air-flow 73.63 m3/h, SPL = 5.71 dB/A, NB-eLoop_®_ 140mm, Blacknoise, Germany), for air in- and outlet.
- A recording camera (GigE Basler ½” NIR, NOLDUS, Germany), equipped with a varifocal lens (4.5 – 12.5 mm, NOLDUS, Germany) and infrared pass filter.

### Pilot studies

We performed a series of small studies in a cohort of 10 animals (5 animals under up-phase and 5 under down-phase CLAS), to investigate the most adequate set of stimulation parameters. Briefly, we opted to stimulate at 35 dB (range tested: 50 dB, 45 dB, 40 dB and 35 dB) based on real-time video and posterior EEG/EMG inspection, as none of the animals showed any perturbation during sustained NREM sleep under this volume. Based on literature (Fattinger et al., 2017; H.-Viet V. Ngo et al., 2013), and considering average delay from detection to trigger of 29.30 ms, we selected 60° for up-phase and 180° for down-phase stimulation (we defined slow wave’s 0° as the rising zero-crossing, 90° to the positive peak and 270° to the slow-wave trough). Phase-target CLAS in humans typically uses 50 ms stimulus duration for a dominant slow oscillatory component of 0.8 Hz. Therefore, we determined 30 ms of stimulus duration to overlap with the ongoing slow waves proportionally — slow oscillatory activity peaks at 1.35 Hz in rats (Mölle, Eschenko, Gais, Sara, & Born, 2009).

### Experimental design

We performed the whole experiment in four batches of 5 or 6 animals each, divided evenly across experimental groups. Aiming to test several subjects daily, individually and at the same circadian time, the light/dark cycle of each subject was adapted accordingly. Briefly, we shifted the light/dark cycle at once, allocating the animals’ first hour of light into successive testing windows of 1.5 h during the course of the working day. Bearing in mind that, during the testing period, the circadian cycle of the last animal of each batch was shifted 9 hours in relation to the first, the entire batch was given a 10-day adaptation to the new routine, as well as to the recording chamber and cables. We kept the animals on a feeding schedule in which they daily received 50 g/kg of regular chow after training throughout the entire experiment. This feeding schedule did not cause any significant weight loss. Water was available ad libitum. Following 24 hours of baseline recording (BL), we initiated a 16-day auditory stimulation protocol in parallel with a fine motor-skill learning task (see *Single pellet-reaching task protocol*). On the 17^th^ protocol-day, we performed the last behavioral assessment and sacrificed the animals (**Fig. 2**).

### EEG/EMG recording and pre-processing

In order to verify the effect of auditory stimulation on EEG spectra, we conducted bilateral tethered EEG/EMG recordings (differential mode) during 24 h, to serve as BL, and throughout all the subsequent 16 protocol-days, applying our runtime stimulation paradigm in 5 or 6 freely-moving animals simultaneously. We acquired data using a Multichannel Neurophysiology Recording System (Tucker Davis Technologies, TDT, USA). We sampled all EEG/EMG signal at 610.35 Hz, amplified (PZ5 NeuroDigitizer preamplifier, TDT, USA) after applying an anti-aliasing low-pass filter (45% of sampling frequency), synchronously digitized (RZ2 BIOAMP processor, TDT, USA), recorded using SYNAPSE software (TDT, USA) and stored locally (WS-8 workstation, TDT, USA). We filtered real-time EEG between 0.1 – 36.0 Hz (2^nd^ order biquad filter, TDT, USA), and EMG between 5.0 – 525.0 Hz (2^nd^ order biquad filter and 40-dB notch filter centered at 50 Hz, TDT, USA), and feed the signals to real-time detection algorithms for NREM staging and phase detection (**Fig. 1c**).

### Online NREM staging and auditory closed loop stimulation

Parallel rule-based NREM staging and phase detection features were continuously running alongside EEG/EMG recording, i.e. sound triggers were presented at every instance the stimulatory truth function compounding these features was reached (**Fig. 1c** and **Fig. 1-figure supplement 1**). For online sleep staging, a non-linear classifier compounds two major decision nodes: power in EEG and power in EMG (Hamrahi, Chan, & Horner, 2001). Briefly, we computed high-beta (20 – 30 Hz) and delta (0.5 – 4 Hz) bands’ rms on a sliding window of 1 s, using an algorithm written in RPvdsEx (Real-time Processor visual design studio, TDT, USA). Once the ratio rms_delta_/rms_high-beta_, hereinafter referred as NREMratio, crossed a threshold individually identified during the BL recording, we further confronted EMG rms to a threshold, as well defined during BL, in order to rule out movement artifacts. The above-mentioned NREMratio and the EMG power thresholds for online NREM staging during auditory stimulation, suggestive of sustained SWS during NREM, were extracted individually and immediately after the BL recording: 24-h EEG/EMG data was automatically scored (SPINDLE, ETH Zurich, Switzerland) and fed to a custom-written MATLAB (ver. R2016b) script. In short, a strict estimate of NREMratio threshold during NREM was established as +1.0 SD over the mean (representing the 84.1% percentile) of the NREMratio of all NREM epochs during BL. Similarly, EMG power in NREM was delimited to values −1.0 SD below the mean (threshold at 15.9% percentile) of the EMG rms values during offline-scored NREM sleep. These two values marked the transition into consolidated NREM sleep in each subject during online NREM staging, and were introduced to our customized SYNAPSE_®_ (TDT, USA) project. For phase-targeted auditory stimulation of slow waves, SYNAPSE_®_ combines the online NREM-staging feature with a phase-detector. Briefly, a runtime very-narrow bandpass filter (TDT, USA) for EEG phase detection isolated the 1-Hz component for phase targeting of slow waves (approximately 1.35 Hz in rats (Mölle et al., 2009)) of each subjects’ left EEG channel. We predetermined slow wave’s 0° as the rising zero crossing, 90° to the positive peak and 270° to the slow-wave trough. At every identified positive zero-crossing on the filtered signal, the phase detector resets to 0°, and calculates any of the selected target phases based on the number of elapsed samples since the zero crossing. This method offers the chance to recognize slow waves consistently across conditions, independently of the target-phase. We divided the animals into three different phase-targeted stimulation approaches: up-phase stimulation targeting 60°, down-phase stimulation targeting 180°, and mock stimulation flagging slow waves at 0° (arbitrarily chosen) with no delivery of sound (**Fig. 1d**). On average, a sound trigger was sent 29.30 ms (RX8 MULTI-I/O processor, TDT) upon validation of truth-value for NREMratio, EMG power and phase target criteria. Auditory stimuli consisted of clicks of pink 1/f noise (30 ms duration, 35 dB SPL, 2 ms rising and falling slopes) in free field conditions, from built-in speakers (MF1 Multi-field magnetic speakers, TDT) on top of the stimulation chamber, 50 cm above the center of the floor-area). The protocol of the mock condition was identical to up-phase or down-phase stimulations, but the sound was muted. All triggers were time-flagged for offline analysis. Throughout all days of stimulation, we used a video system to sporadically control for stimuli-evoked arousals or reflexes.

### EEG/EMG scoring

We scored all recording files using the online computational tool SPINDLE (Sleep phase identification with neural networks for domain-invariant learning) (Miladinović et al., 2019) for animal sleep data. In short, European Data Format (.edf) files, consisting of two parietal EEG and one nuchal EMG channels, were uploaded to SPINDLE to retrieved vigilance states with 4-second epoch resolution. The algorithm classified three vigilance states: wakefulness, NREM sleep, and REM sleep. Additionally, unclear epochs or interfering signals were labeled as artifacts in wakefulness, NREM, and REM sleep. Wakefulness was defined based on high or phasic EMG activity for more than 50% of the epoch duration and low amplitude but high frequency EEG. NREM sleep was characterized by reduced or no EMG activity, increased EEG power in the frequency band < 4 Hz, and the presence of SOs. REM sleep was defined based on high theta power (6 – 9 Hz frequency band) and low muscle tone.

### Post-processing of EEG

Time spent in NREM was determined as an absolute number of minutes for BL, 3rd day of motor training (M-T_3_), and last day of non-training period (N-T_4_), representing an unperturbed (no stimulation) day before the stimulation runtime, the last stimulated day of motor-training protocol (11 days after BL), and the last stimulated day of the designed protocol (16 days after BL), respectively. For the same days, we also calculated a NREM fragmentation index, expressed as the number of NREM bouts/total number of NREM epochs, with higher index values expressing higher fragmentation. Furthermore, we extracted measures of global spectral responses in specific bandwidths, by processing the left-hemisphere EEG signal with a custom MATLAB routine (ver. R2016b). Briefly, the EEG signal was resampled at 300 Hz, and multiplied with a basic Fermi window function *f*(*n*) = (1 + _e_^(5-*n*/50)^)^(−1)^, to gradually attenuate the first and last 2 s (n = 600). Next, we filtered the signal between 0.5 - 48 Hz using low- and high-pass zero-phased equiripple FIR filters (Parks-McClellan algorithm; applied in both directions (*filtfilt*); order_high_ = 1880, orders_low_ = 398; −6 dB (half-amplitude) cutoff: high-pass = 0.28 Hz, low-pass = 49.12 Hz). The signal was visually inspected for regional artifacts (2-hour sliding window) not detected during automatic scoring: within scored NREM sleep, brief portions (< 10 sample-points at 300 Hz) of signal > ± 8 × interquartile range were reconstructed by piecewise cubic spline interpolation from neighboring points. We performed spectral analysis of consecutive 4-s epochs (FFT routine, Hamming-window, 2 s overlap, resolution of 0.25 Hz), and normalized the power estimate of each frequency-bin in relation to the total spectral power (0.5 – 30 Hz). Additionally, we calculated the hourly activity in each frequency-band — delta (>0.5 – 4 Hz), theta (>4 – 8 Hz), alpha (>8 – 11 Hz), sigma (>11 – 16 Hz), and beta (>16 – 20 Hz) — as the hourly mean power of NREM epochs, using the abovementioned digital filters (order_high_ = 3758 and orders_low_ = 3861), normalized by the hourly total power (0.5 – 30 Hz). Relative 24 h delta- and sigma-power were calculated in a similar fashion for M-T days and subsequently grand-averaged for that period.

### Post-processing of sound triggers and evoked-response potentials

We analyzed the trigger distribution during all stimulation days in 3 aspects: the precision of stimulation as the ratio between the number of triggers during NREM sleep and the total number of triggers; trains of triggers as the count of any size sequences of triggers 1 second or less apart (as SOs in rats occur at a peak frequency of 1.35 Hz)(Mölle et al., 2009); and interstimulus interval (ISI) as the time between triggers (**Fig. 1e**). Analysis of the evoked auditory event-related potentials (ERPs) was performed computing EEG snippets across all auditory triggers in NREM, within a 3.5 s window time-locked to the trigger onset, and a post-stimulus cut-off of 2 s. Only triggers with a post-ISI > 1 s were included in this analysis. To assess the ability of the runtime paradigm to target 60° (up-phase) and 180° (down-phase), we filtered the left EEG channel between 0.5 and 2 Hz (same fashion as previous digital filters: order_high_ = 3758, orders_low_ = 3861; −6 dB (half-amplitude) cutoff: high-pass = 0.39 Hz, low-pass = 1.91 Hz), followed by a Hilbert transform to define instantaneous phase. We averaged the instantaneous phase for all triggers delivered in NREM during the 16-day stimulation period and plotted the results in histograms with 6 bins of 60°. Counts were normalized to the total number of counts in each animal. Summary statistics were computed using the CircStats toolbox in Matlab (Berens, 2009).

### Single pellet-reaching task protocol

We used a 15 × 40 × 30 cm (W × L × H) acrylic-glass chamber with a vertical window (1 cm wide, lower edge 5 cm above the cage floor) in the front wall. This window was covered by a motorized sliding door, connected to an inductive sensor in the back wall. The rat could open the mentioned door by poking the nose to the sensor in the back wall of the cage. The training procedure consisted of 2 phases: 1) habituation to the testing chamber and instrumental- or pre-training during which the rat is taught to operate the motorized door (further called P-T), and 2) reaching- or motor-training during which the rat is taught to use a single paw to obtain food pellets (further called M-T) (**Fig. 1f**). In detail:

1. P-T: We food-restricted the animals for 24 h before the first day of the protocol to ensure engagement in the task. We introduced all animals to the testing chamber for 3 days, during 60 minutes daily, and allowed familiarization to the cage and the positioning of the food pellets (45 mg, Bioserve Inc., Frenchtown, USA). On the 4^th^ day and for 4 days, we started the pre-training routine by guiding all animals to the nose-poke sensor in the back of the cage, enabling the motorized door covering the front window to open. This routine was rewarded with a single food pellet placed at the window edge, easily reached using the tongue. Each P-T session consisted of 100 trials - opening the window and retrieving a pellet - or 60 minutes, whichever occurred first.
2. M-T: Subsequent skilled reaching training was similar to P-T, but we placed the pellet on a pedestal (7 mm diameter), located 1.5 cm outside of the cage. In this position, the rat could only use a forelimb to retrieve the pellet (**Fig. 1g**). On the first day, each animal quickly (during the first 5-10 reaches) demonstrated a forelimb preference, determining the placement side of the pedestal (left side if right forelimb was preferred and vice versa). The right forelimb was preferred in 5 subjects in the up-phase group (n=8), in 6 in the down-phase group (n=8), and 2 in the mock group (n=6). In addition, in the mock group, 2 subjects showed equal preference for left and right grasping, to which we placed the pedestal on the left. Overall, we rarely saw the animals using the non-preferred forelimb. A trial ended when the pellet was grasped or pushed off the pedestal, automatically closing the door within 1 s (via built-in inductive sensor on the pedestal that sensed the presence of the pellet). We scored a reaching attempt, or occasionally the last of multiple reaching attempts per pellet, as either successful (pellet is retrieved from the pedestal and eaten), failed (pellet is pushed off the pedestal) or drop-in (rat grasps the pellet but drops it during paw retraction outside or inside the cage). Each M-T session consisted of 100 attempts or 60 minutes, whichever occurred first, divided into 5 bins of 20 pellets or 12 minutes (**Fig. 1h**).

### Single pellet-reaching task analysis

We defined success rate (SR) as the number of successes out of 100 possible attempts. We calculated the intra-session change as the log2 of the fold-change of the number of successes of the last bin to the first bin of a session (−1 represents a 50% drop in successes, while 1 represents twice more successes from first bin). Rats that failed to achieve a minimum SR of 10% at M-T_2_ or after would be excluded, but no subject met this criteria.

### Statistical analysis

We present all data as the mean ± SEM, unless stated otherwise. We performed statistical analyses using Prism 8.0 (GraphPad Inc., San Diego, CA, USA), SPSS Statistics 26.0 (IBM Corp., Armonk, NY, United States), R 4.0.2 (Boston, MA, United States)(Bates, Mächler, Bolker, & Walker, 2015; Lenth, 2020; Team, 2020), and Matlab R2016b and R2019a (Natick, MA, USA). We performed power calculations (R 4.0.2, *power.anova.test*) based on human literature using the same stimulation paradigms. We expected a standard deviation of 15% for all group estimates referent to delta activity change from baseline (up-phase = +15% (H.-Viet V. Ngo et al., 2013; Papalambros et al., 2017); mock = 0%; down-phase = −10% (Fattinger et al., 2017; H.-Viet V. Ngo et al., 2013)), to which we obtained a sample size n = 8 (for sigma level = 0.05 and power = 0.80). Multiple two-way repeated measures analysis of variance (RM-ANOVA, adjusted using the Greenhouse-Geisser correction when necessary) were used to assess the influence of *time* (days, or bins in the SPRT analysis) and *condition* (up-phase *vs* mock, and down-phase *vs* mock) on NREM proportions, time-course of delta and sigma, and SR on last protocol-day. Paired Wilcoxon rank tests were applied to NREM amount relative to BL values, since data did not follow a normal distribution for the plotted protocol-days. We used a one-sample Hotelling’s T2 test to explore whether the daily changes in all frequency bands has a mean equal to a null change (Oja, 2010). Differences on the time-course of delta- and sigma- frequencies in relation to BL were tested using multiple paired t-tests, corrected with Holm-Sidak method for multiple comparison. Phase targeting was evaluated using circular statistics from the CircStat toolbox(Berens, 2009). For the analysis of SPRT counts, Poisson distribution or, if dispersion was larger than expected, Negative Binomial distribution was fit to the data. Both models provide Incidence Rate Ratios (IRR) for which the expected counts were estimated and analyzed using mixed regression models (fixed factors: days and groups; random factor: subjects) followed Tukey’s multiple comparison tests. Significance for the models’ fixed effects was estimated with Type II Wald chi-square tests. Associations were tested using 1-tailed Spearman’s rank correlations, underlying established one-directional hypotheses. Unless stated otherwise: *n* (up-phase) = 7; *n* (mock) = 5 for electrophysiology results and *n* (mock) = 6 for behavioral results; *n* (down-phase) = 8. The alpha level for all statistical tests was set initially to 0.05.

## Contributions

CRB and DN designed the study. CGM executed the experiments. CGM, MS, SIN, and SL curated and analyzed the data. CGM, CRB, RH, and DN interpreted the results and wrote the manuscript. All authors corrected and approved the final version of the manuscript.

## Data availability

The raw data blocks that support the development of this tool are available upon request to the corresponding author. All figures accompanied by an Excel file containing the numerical data and statistical analyses are provided with this submission.

## Acknowledgements

We thank Mr. Mark Hanus, Mr. Myles Billard, Dr. Clément Vitrac and Laura Ferster for their valuable support.

---

**Figure 1-figure supplement 1.**
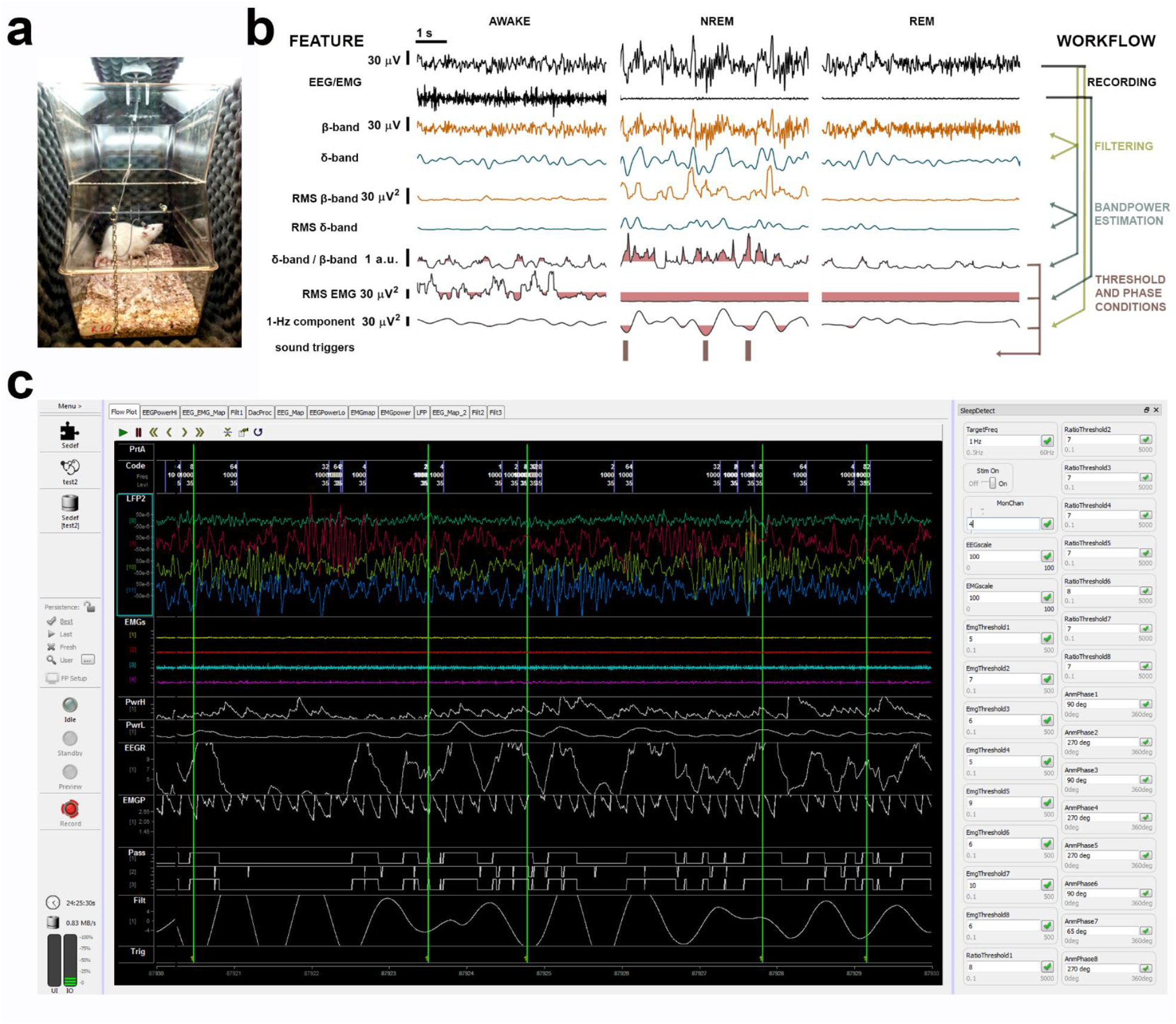
**Illustration of a sound insulated chamber, online signal-processing workflow and SYNAPSE’s interface. a.** Single sound insulated chamber for recording and auditory stimulation in a freely moving rat. **b.** Processing tree for online NREM-staging and phase-targeted auditory stimulation: left EEG and EMG channels are filtered and analysed for spectral power. Based on individual thresholds predefined during BL (highlighted pink areas for δ-band/β-band (i.e. NREMratio) and EMGrms), auditory triggers were delivered in selected phases of slow waves, during online-staged NREM. **c.** Synapse interface depicting EEG/EMG signal (in colour) and phase-locked stimulation triggers (vertical green lines) in 4 subjects. On the right, input boxes for individual NREMratio and EMGrms thresholds and phase-targets will determine the optimal timing for phase-locked auditory stimulation during NREM in each animal.

**Figure 5-figure supplement 1.**
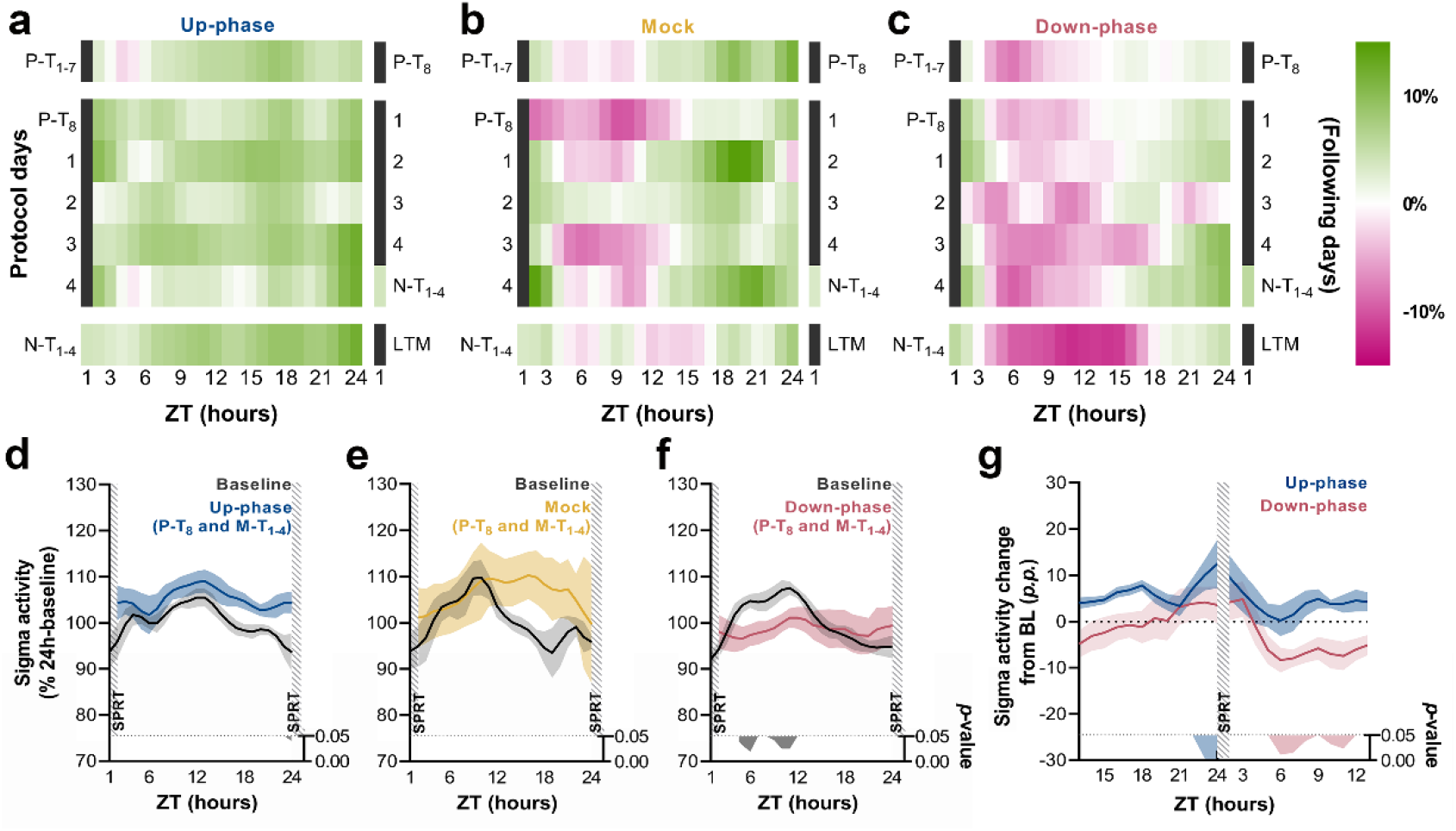
Time-course of sigma activity changes from baseline during all protocol days and motor-training grand-average. **a-c.** Sigma activity (11 – 16 Hz) changes from baseline, in 1-h bins, for up-phase, mock, and down-phase stimulated animals. P-T and N-T days were condensed into grand averages (top and bottom rows, respectively). Individual M-T days are shown in the middle rows. Hours in black represent the single-pellet reaching task training sessions, numbered with the respective protocol-day. On the right side of each heat-map, the training sessions of the following protocol-day are enumerated. **d-f.**-Time-course of hourly sigma-activity during BL and motor-training days, collectively (coloured line represents the grand-average of training sessions P-T_8_ to M-T_4_). **g.** Contrast between delta activity during the single-pellet reaching task session’s preceding and succeeding hours for up- and down-phase stimulation groups, plotted in percentage points from BL. Grey/coloured shadows bellow plots D-G represent significant time-points (multiple t-tests, Holm-Sidak correction for multiple comparisons). ZT: zeitgeber. P-T: pre-training. M-T: motor-training. N-T: no-training. LTM: long-term memory. SPRT: single-pellet reaching task. p.p. percentage points.

**Figure 6-figure supplement 1.**
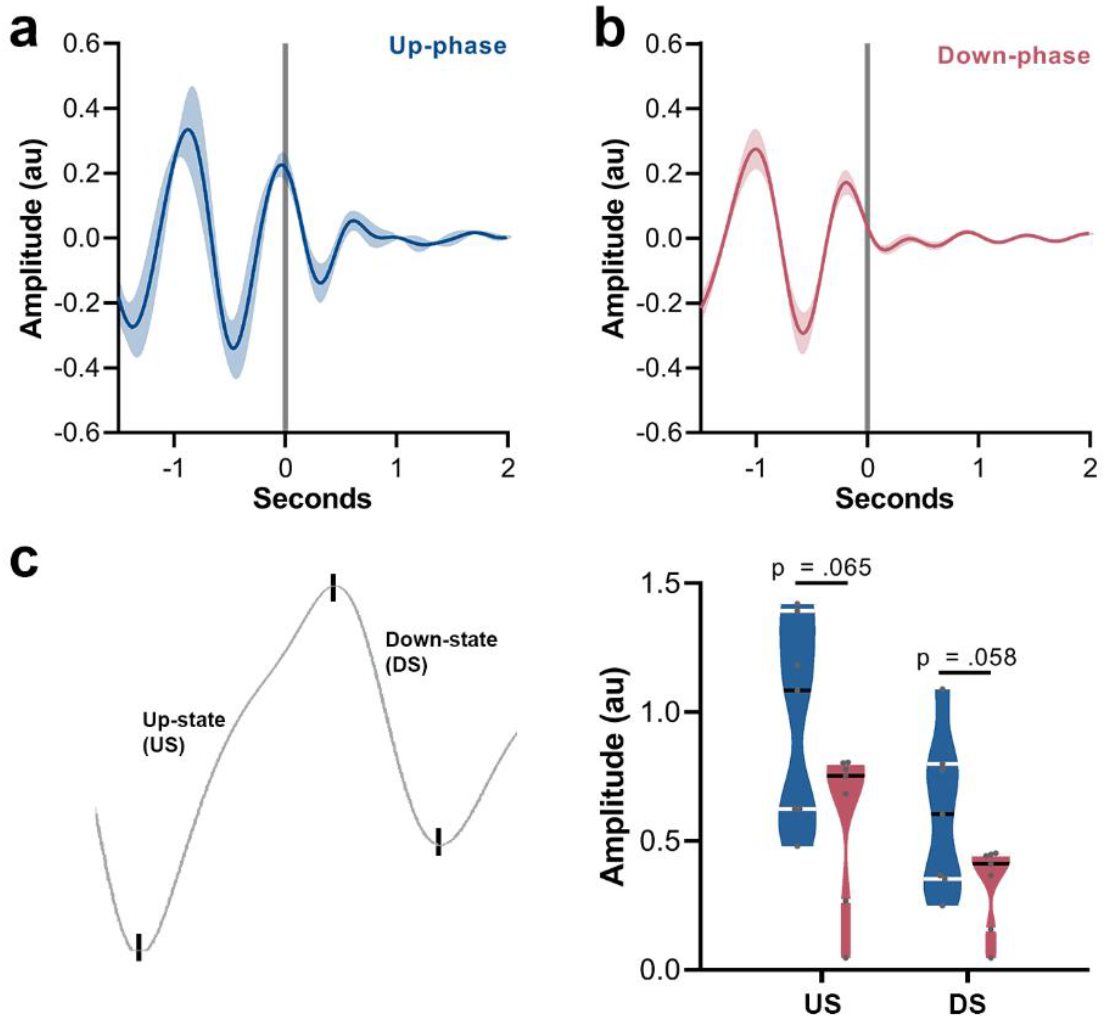
Grand mean ERP of up- and down-phase groups, and amplitude contrast between the two conditions. **a.** Waveforms from M-T period were locked to the start of each auditory trigger onset (t = 0). Up-phase seems to induce a second slow wave cycle. **b.** Down-phase stimulation does not preserve the slow oscillatory pattern. **c.** Comparison of amplitude between up- and down-phase conditions for the depolarized up-state (unpaired t-test, *p* = .065) and the hyperpolarized down-state (unpaired t-test, *p* = .058). US: up-state. DS: down-state. au: arbitrary units.

